# CIAdex: Single-Cell FTIR Spectral Fingerprinting for Cell Identity Verification and Aging Quantification in Therapeutic Cell Manufacturing

**DOI:** 10.1101/2025.07.31.667910

**Authors:** Ying Wang, Yanfei Ding, Chengzhang He, Xiaojie Zhou, Junwu Tu, Yulei Deng, Jian Zhao

## Abstract

Ensuring the identity and optimal aging state of cell products is critical for the efficacy and safety of cell therapies. Despite rapid iterations, there remains an urgent need for robust and easy-to-implement tests to characterize cell products. Here, we present CIAdex (<u>C</u>ell Identification and <u>A</u>ging In<u>dex</u>), an analytical framework that utilizes single-cell Fourier-transform infrared (FTIR) spectral fingerprints and machine learning to achieve precise label-free cell identity assessment and aging quantification. CIAdex employs a Feature Extraction Processor to automatically extract FTIR spectral variables corresponding to distinct cellular biomolecular features, enabling reliable distinction of lineage-, donor-, and batch-specific cell populations using linear discriminant analysis. Through application of the XGBoost algorithm, a quantitative aging index (trPDL/trPN) was further generated for tracking cellular aging dynamics along culture expansion. Notably, trPDL/trPN quantitatively represent subtle age-related shifts among different batches and drug-induced senescence or rejuvenation effects which are unmeasurable by existing methods. Together, our work demonstrates that CIAdex, by simultaneous label-free identity verification and aging quantification of cell populations, offers a transformative approach to interpret single-cell FTIR spectral fingerprints and provides novel metrics for quality control in cell manufacturing with significant potential for optimization and assurance of cell therapies’ safety and efficacy.

## Introduction

Cell-based products ranging from pluripotent stem cells, lineage-specific progenitor cells, to function-defined stromal cells and somatic cells hold immense promise for repairing, replacing, or regenerating tissues and organs against neurodegenerative disorders, cardiovascular diseases, wound healing, etc. The promise of cell therapies, especially stem cell therapies, is based on the inherent biological properties of living cells, including proliferation/self-renewal, plasticity, migration/integration, etc. However, donor variability, batch-to-batch inconsistencies, and subtle culture variations introduce significant heterogeneity in cell products, which pose particular challenges for product development as well as compromise the safety and efficacy of these therapies^1–7^. Crucially, cellular aging due to both the chronological age of the donor and the cell aging process during *in vitro* expansion increases the risk of chromosomal abnormalities or mutations and accumulates epigenetic alterations in cells, which affect gene expression profiles and directly impact the cellular function and the therapeutic potential. For instance, mesenchymal stromal cells (MSCs) exhibit diminished proliferation and immunomodulatory capacity after prolonged *in vitro* expansion or from elderly donors^8^. Neural progenitor cells (NPCs) lose proliferation and neuronal differentiation potential during extended culture^9^. Bone marrow-derived stromal cells from young patients demonstrate superior skin healing efficiency in either autologous or allogeneic transplantation compared to those from older patients^10^.

Advances in cellular therapy and regenerative medicine have accelerated the development of quality assurance and control measurements, including cell identity verification, assessments of cell viability, purity, genomic stability, senescent status, functional potency, and other metrics. Nevertheless, existing approaches, such as immunofluorescent imaging, flow cytometry, single-cell RNA sequencing, and other multi-omics analysis, are destructive, low-throughput, labour-intensive, time-consuming, and often difficult for real-time capturing of cellular and molecular heterogeneity and dynamics^14–18^. The cumulative population doubling levels (cPDLs) have been widely recommended for tracking cellular aging in addition to the passage numbers^19^. However, cPDLs still rely on manual record-keeping and could become unreliable under variable culture conditions or drug treatments; therefore, they are problematic for standardization across institutions. On the other hand, to mitigate the effects of aging on cell therapy outcomes, senescence markers (e.g., β-galactosidase, p16, etc.) have been commonly applied for aging status assessment, but only at static endpoints, as they lack sensitivity to track progressive aging phenotypes. Moreover, structural variations (e.g., lipid unsaturation, chromatin conformation) that precede functional decline have not yet been explored and quantified for their potential as an early aging index. While regulatory agencies, including ICH, FDA, EMA, etc., emphasize rigorous quality control and assurance of cell-based products^11–13^, we still lack robust “snapshot” tools to directly verify cell identity and track molecular changes linked to donor, batch effects, and expansion variation that may impact therapeutic efficacy. These challenges underscore the urgent need for a robust, label-free, and unified in-process analysis to resolve cell population identification and verification, batch consistency assessment, and aging measurement beyond cPDL/passage counting.

Fourier transform infrared (FTIR) microspectroscopy emerges as a promising solution, providing non-destructive, label-free, high-resolution molecular fingerprinting of single cells^20^. By capturing vibrational signatures of biomolecules (e.g., lipids, proteins, nucleic acids), FTIR offers insights into both cellular compositional and structural differences that are crucial for assessing cell quality^10,21^. Prior studies demonstrate the utility of FTIR enhanced by synchrotron radiation (SR-FTIR) in distinguishing cell types and identifying pathological states^22–26^. However, its application in quality assurance and control or in cell aging dynamic monitoring for cell manufacturing remains underexplored.

We hypothesize that FTIR spectral features, reflecting the unique biomolecular profiles of cells, can be leveraged to assess cell identity and aging status. Our work indeed shows that, by integrating machine learning, we can unlock these FTIR spectral data for both cell population identity and aging dynamics, thereby offering new quality control metrics for cell product manufacture as well as efficacy validation and optimization of cell therapies.

## Results

### Discrimination of cell populations with single-cell FTIR spectral fingerprints across cell types, donors, and batches

FTIR spectroscopy detects vibrations of biological components and thereby provides high-resolution, label-free cellular molecular fingerprints. First, we test whether FTIR microspectroscopy can be applied to distinguish different cell types from the same donor. Human iNPCs (induced neural progenitor cells) were derived from the human induced pluripotent stem cell (iPSC) line TUSMi002-A generated from a healthy female^29^. The characteristics and purity of the TUSMi002-A-iPSCs and their derivatives, TUSMi002-A-iNPCs, were assessed (Figure S1). Then, TUSMi002-A-iPSCs and TUSMi002-A-iNPCs were cultured on calcium fluoride (CaF_2_) slides and analyzed via SR-FTIR microspectroscopy to obtain high-resolution single-cell spectra. The infrared images of individual iPSCs and iNPCs are shown in Figure 1a. The biochemical heatmap images display distinct distributions of biomolecules, including fatty acids, proteins, and nucleic acids in each cell, where red indicates higher concentration and blue indicates lower concentration. These cellular components exhibited unique spatial intensity patterns between the two cell types from the same donor. The raw single-cell FTIR absorbance spectra and their second derivatives for individual iPSCs and iNPCs are shown in Figure 1b. The spectral second derivatives show distinct peaks across wavenumbers from 3400 to 1000 cm□¹, with notable differences between iPSCs and their iNPC derivatives, suggesting FTIR spectra could be used to represent the distinct molecular compositions between the two cell types. Then, the second derivatives of the original absorbance bands were applied for the quantification of 27 specific spectral absorption peaks, which correspond to distinct chemical functional groups of intracellular compositions and structural elements as reported previously^25,31–33^ (Figure S2). We first developed a gradient-boosted decision tree-based feature extraction processor (FEP) for automatic and unbiased extraction of these 27 feature peaks. FEP functions to determine appropriate wavenumber ranges around target characteristic peaks with factors including peak order, morphology, and overall spectral distribution patterns. Then the 27 intrinsic FTIR spectral variables of TUSMi002-A iPSCs and derived iNPCs were extracted and subjected to linear discriminant analysis (LDA), the supervised dimensionality reduction technique^34^, for separating iPSC and iNPC populations. As shown in Figure 1c, LDA plots demonstrated clear segregation, with iPSCs (black) and iNPCs (red) forming non-overlapping frequency distributions. The corresponding F-score plot shows key characteristic peaks (numbered 3, 7, 19, 25, and 27) corresponding to the energy of molecular vibrations for Olefinic =CH stretching, Vs(CH_2_), CH_3_ bending and CH_2_ scissoring vibrations, DNA C-O stretching vibrations and glycogen as the most significant discriminators. Interestingly, critical differences arose not from absolute absorption intensities that reflected the amount of infrared light absorbed at each functional group but from shifts in vibrational energy levels reflecting structural variations (e.g., lipid unsaturation, DNA conformation). These results indicate that FTIR microspectroscopy captures the differences in molecular compositional and structural fingerprints between iPSCs and iNPCs, suggesting that high-dimensional FTIR spectral data could be applied for the discrimination of different cell types with the most distinct cellular biochemical features highlighted.

**Figure 1.**
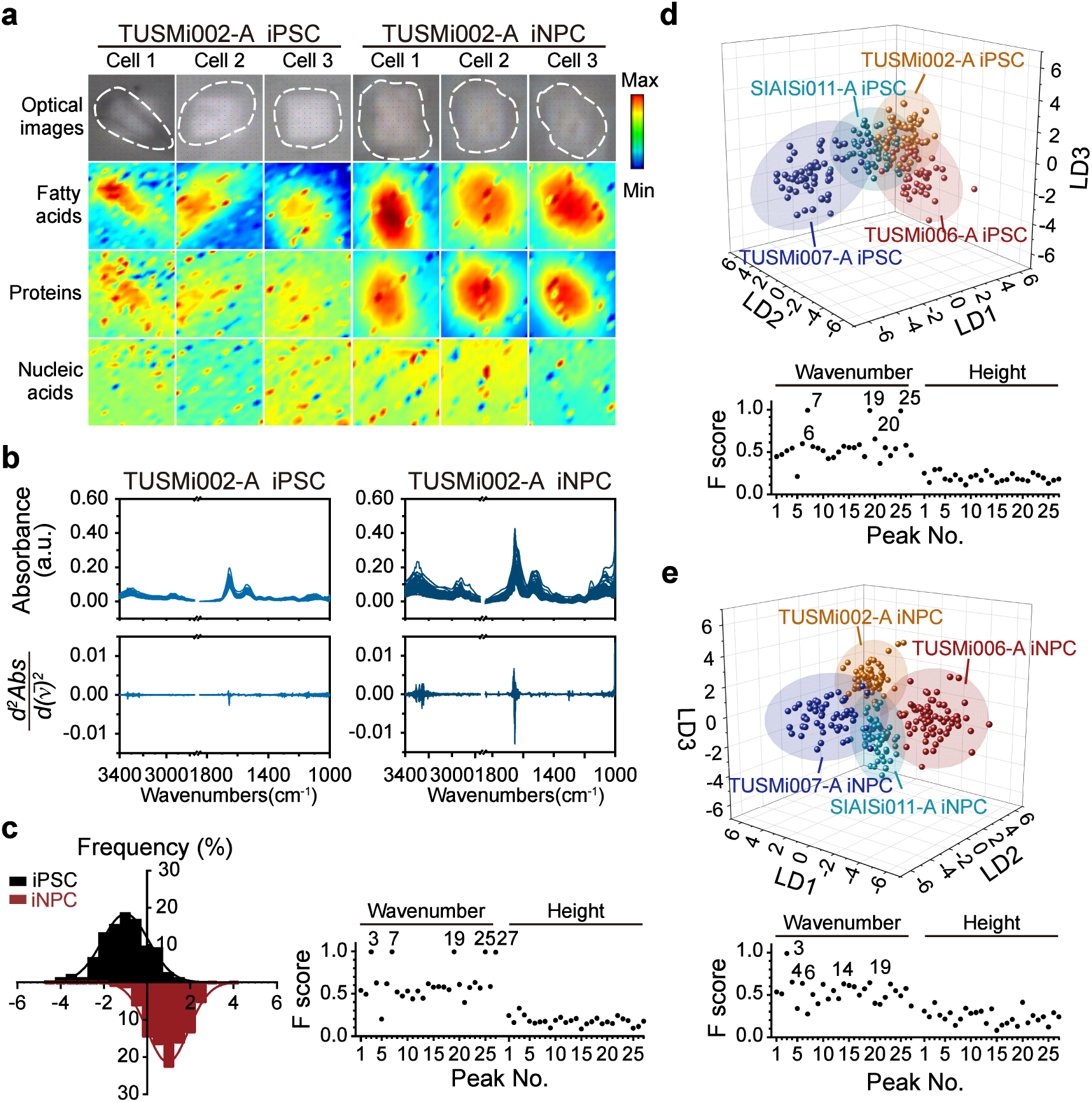
Single-cell FTIR spectral fingerprints decode cell identities. **a.** Single-cell FTIR microspectroscopy. Representative bright field images and infrared heatmap images of individual iPSCs or iNPCs were shown. The infrared heatmap images exhibit a distinct distribution of fatty acids, proteins, and nucleic acids in each cell, based on spatial intensity patterns in 3000-2830 cm^−1^ (fatty acid), 1770-1475 cm^−1^ (protein), and 1300-1000 cm^−1^ (nucleic acid), respectively. The colour scale indicates intensity of absorption corresponding to concentration of biochemical molecules from lowest (blue) to highest (red). **b.** The raw FTIR absorbance spectra measurements (top graphs) and their second derivatives (bottom graphs) for individual TUSMi002-A iPSC and iNPC cells. Both are plotted against wavenumbers (cm^−1^) ranging from 1000 to 3400. **c.** LDA frequency distribution of TUSMi002-A iPSC (black) and derived iNPC (red) on 2D plot (left). F-scores for the wavenumber and height of each peak were plotted, and the peak numbers for the five features with the highest F-scores were shown (right). **d-e**. 3D scatter plots of LDA cluster analysis of iPSCs (d) or iNPCs (e) from different donors (TUSMi002-A, TUSMi006-A, TUSMi007-A, and SIAISi011-A). The top plots showed a clear separation between different donor-specific cell populations. The bottom plot shows F-scores for the wavenumber and height of each peak. The peak numbers for the five features with the highest F-scores were shown.

Then we asked whether we can apply FTIR microspectroscopy for distinguishing cell populations of the same cell type but from different donors. Conventionally, the verification of cell lines from different donors can only be achieved using STR analysis. The characteristics and purity of iPSCs and iNPCs derived from different donors (TUSMi006-A^35^, TUSMi007-A^36^, SIAISi011-A^37^) were assessed (Figure S1). LDA of the 27 variables resolved donor-specific iPSCs into distinct clusters (Figure 1d). Similarly, iNPCs from different donors formed separate clusters in 3D LDA space (Figure 1e). These results confirm the capability of our analytical strategy for resolving cell-type-specific and donor-specific variations, with peaks 3, 7, 19, and 25 remaining critical discriminators. Notably, donor-specific differences in peak 25 (DNA C-O stretching vibration) aligned with minor epigenetic variability, suggesting that spectral fingerprints may reflect subtle molecular heterogeneity.

Then we tested whether these 27 intrinsic absorption peaks enable precise distinguishing between different cell batches of individual cell lines. To test this, we implemented LDA on the second derivative FTIR spectra of different batches from TUSMi002-A iNPCs (labeled as TUSMi002-A-1 to TUSMi002-A-5) and compared these various batches with SIAISi011-A iNPCs (Figure S3a), TUSMi006-A iNPCs (Figure S3b), or TUSMi007-A iNPCs (Figure S3c). The LDA plots show well-defined clustering patterns with the various batches of TUSMi002-A iNPCs, which were also well separated from SIAISi011-A iNPCs, TUSMi006-A iNPCs, or TUSMi007-A iNPCs, validating the robustness of the FTIR microspectroscopy-based analytical framework for distinguishing inter-batch variations. Interestingly, the F-score analysis for these comparisons reveals consistent discriminative molecular features, including amide N-H bending, stretching vibrations of glycogen, and nucleic acid ribose linked with DNA breaks and PARylation^38^, which are particularly important in distinguishing between different cell batches.

Together, these results indicate that this FTIR microspectroscopy-based analytical framework not only enables the reliable discerning of cell types, effectively differentiates between various donor-specific cell lines, but also can distinguish among different cell batches of a given cell line based on their molecular fingerprints, providing new attributes for cell population identification and verification.

### Establishment of trPDL/trPN as a quantitative aging index with FTIR Spectral fingerprints

Considering that extended culture can alter the phenotype and function, which further affect the potential therapeutic efficacy of cell products, the cellular age should be carefully tracked during culturing in vitro, and an upper limit needs to be set for the expansion of a given cell population^39^. Based on our observation that different cell batches of a given cell line form clearly-separated clusters in the FTIR spectral analysis with spectral features linked to DNA strand break and PARylation cascades as critical batch-specific discriminators, we hypothesized that FTIR spectroscopy might serve as a reliable, non-invasive tool for characterizing and monitoring cellular aging trajectory through population expansion.

We first recorded population expansion (recorded as cPDLs) and passage numbers of TUSMi002-A iNPCs (Figure 2) and SIAISi011-A iNPCs (Figure S4). As shown in Figure 2a and Figure S4a, we found both TUSMi002-A iNPCs and SIAISi011-A iNPCs exhibited a predictable pattern with cPDL following a quadratic function across cell passages (R² > 0.99), while the population doubling level (PDL) per cell passage, i.e., the population doubling rate, showed a linear decline over passages (R² > 0.89 for TUSMi002-A and R² > 0.85 for SIAISi011-A). Then, by quantitative immunofluorescence analyses (Figure 2b and c for TUSMi002-A, and Figure S4b and c for SIAISi011-A), we found progressive decreases in both proliferation marker Ki67 and key neural progenitor markers Sox2 and Nestin with increasing cPDL/passage number. Cell proliferation analysis (Figure 2d for TUSMi002-A and Figure S4d for SIAISi011-A) confirmed decreased growth rates in higher cPDL populations, which is particularly evident in cells with cPDL65/P25 and cPDL94/P35 compared to early passage cells (cPDL4/P5). We further compared the differentiation potential of iNPCs across various cPDLs (Figure 2e-g for TUSMi002-A, and Figure S4e-g for SIAISi011-A). Immunofluorescence staining for Sox2/Map2 and Tubulin βIII(Tuj1)/Nestin revealed significant differences in neuronal differentiation capacity between low and high cPDL cultures in both cell lines. Quantification (Figure 2f for TUSMi002-A and Figure S4f for SIAISi011-A) showed significant decreases in neuronal markers Map2 (from 68.38% to 28.03% in TUSMi002-A) and Tuj1 (from 66.86% to 31.15% in TUSMi002-A) in high cPDL cells, with similar trends observed in SIAISi011-A. Morphological analysis of differentiated neurons (Figure 2g for TUSMi002-A) revealed progressive reductions in neuronal complexity with increased cPDLs, as evidenced by decreases in branch points (from 3.45 to 0.30), neurite segments (from 8.41 to 1.21), and terminals (from 4.95 to 0.94) in neurons derived from high cPDL TUSMi002-A iNPCs compared with those derived from low cPDL populations. Similar decreases in neuronal complexity were also observed in neurons derived from high cPDL SIAISi011-A iNPCs versus low cPDL population (Figure S4g).

**Figure 2.**
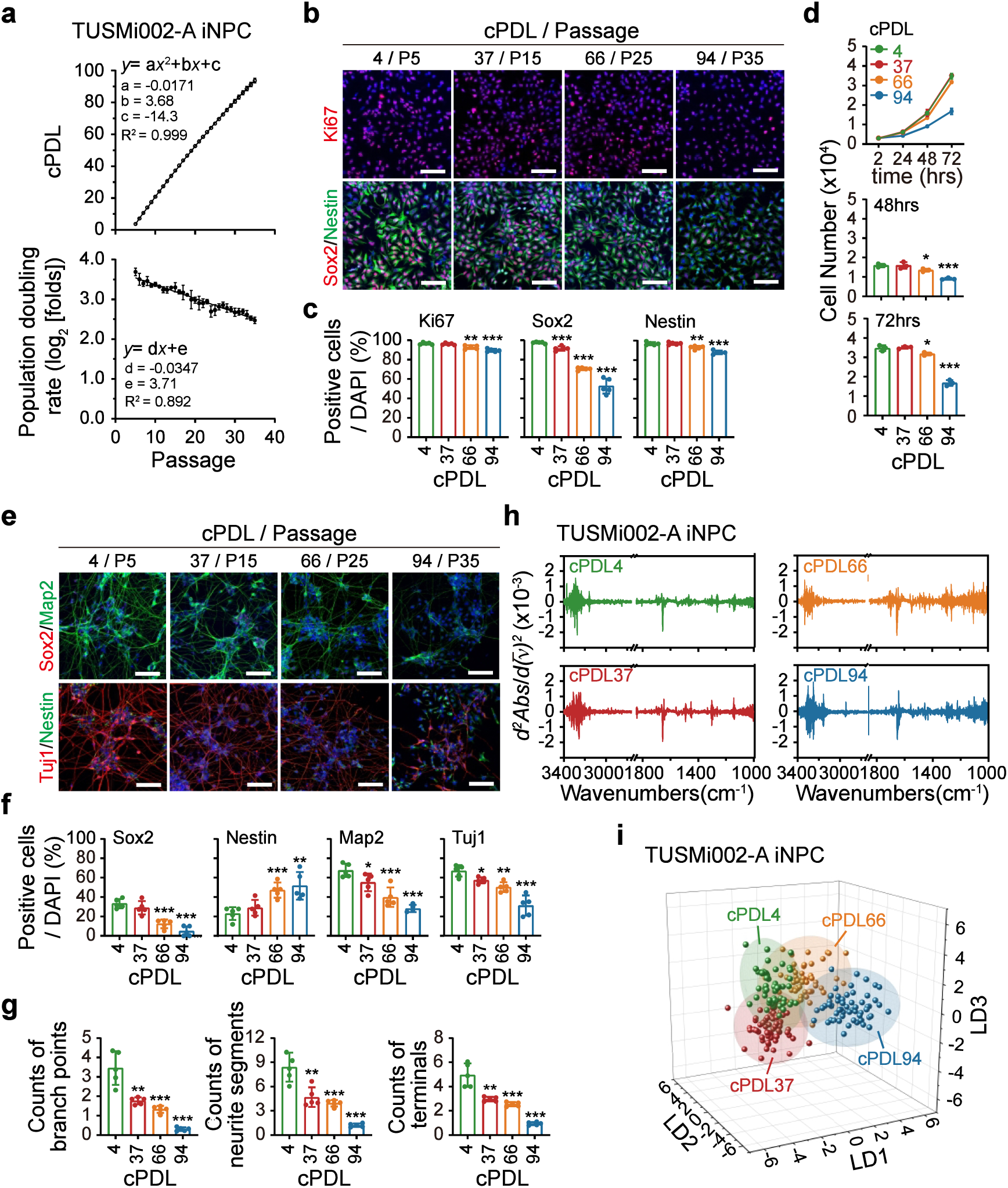
FTIR spectral variables reveal iNPC aging-related biochemical alterations. **a.** The population expansion of TUSMi002-A iNPC over an extended culture period was analyzed and shown. Top: cPDLs (cumulative population doubling levels) were plotted against passage numbers, showing a quadratic relationship (*y* = a*x*² + b*x* + c) with a strong correlation (R² > 0.99). Bottom: population doubling rate (i.e., population doubling level per each culture cycle) decreased as passage number increased, following a linear decay model (*y* = d*x* + e). **b-c**. Immunofluorescent images of Ki67, Sox2, and Nestin in TUSMi002-A iNPCs across four different cPDL/passage points: cPDL4/P5, cPDL37/P15, cPDL66/P25, and cPDL94/P35 (**b**). The percentages for Ki67, Sox2, and Nestin positive cells across the four cPDL points were shown (**c**). Scale bar, 100 µm. **d.** The proliferation of TUSMi002-A iNPCs across the four cPDL points was shown and compared. **e-g**. The differentiation potency of TUSMi002-A iNPCs across the four cPDL points. Cells at the indicated cPDL points were induced to differentiate into neuronal cells. The morphological and immunological characteristics of neuronal cells derived from iNPCs were measured. Map2-positive, Tuj1-positive, Nestin-positive, and Sox2-positive cells in each group were counted and compared **(f)**. Neural morphology was assessed by quantitative analysis of confocal immunofluorescence microscopy images **(g)**. Specifically, the average counts of branches per neuron, neurite segments, and terminals per neuron were compared and shown. Scale bar, 100 µm. **h.** FTIR spectral data and the second derivatives for TUSMi002-A iNPC at different cPDL points were shown. **i.** 3D scatter plots of LDA analysis showed clear clustering of TUSMi002-A iNPC populations across different cPDL points. These results were representative of at least 3 independent experiments. Data were presented as the means ± SEMs. \**P* <0.05, \*\**P* <0.01, and \*\*\**P* <0.001 compared to that of the cPDL4 group. *P* values were determined by a two-tailed T-test.

These results demonstrate systematic phenotypic and functional alterations of iNPCs during extended culture periods, underscoring the importance of monitoring and reporting both cPDLs and passage numbers for standardized cell culture. However, both passage numbers and cPDLs can only be recorded and reported. On the other hand, as shown in Figure 2h-i and Figure S4h-j, we observed that the LDA plots of second derivative FTIR spectra of TUSMi002-A and SIAISi011-A iNPCs showed clear separation between early (PDL4/P5), intermediate (cPDL37/P15), and late (cPDL66/P25 and cPDL94/P35) populations, suggesting that FTIR microspectroscopy could provide an analytical framework for distinguishing molecular signatures associated with different cPDL/passage number cell populations. Therefore, we hypothesized that FTIR spectra might be applied to establish a “transformed” PDL/passage number for quantitatively assessing the cell aging status.

To test this hypothesis, we applied FTIR spectral variables with the correlated cPDLs/passage numbers of different iNPC populations to develop machine-learning models capable of determining the “transformed” PDL/passage number (trPDL/trPN). As shown in Figure 3a, the infrared spectral features of iNPCs with different cPDLs extracted using FEP were constructed as normalized matrices and randomly divided into training and validation data sets. The mean, median, variance, and other statistical metrics of each feature peak in the training set were applied for model development utilizing the XGBoost algorithm. Hyperparameter optimization was performed to determine the optimal learning rate, number of trees, maximum depth of trees, and regularization terms. The validation data set was subsequently fed into the developed model. With the cross-validation strategies accompanied by random sampling combinations, the model’s performance and capacity to generalize across various data sets were confirmed. The calculated single-cell trPDL/trPN values and their associated probabilities with the median and 95% confidence intervals were collected and used as the outputs.

**Figure 3.**
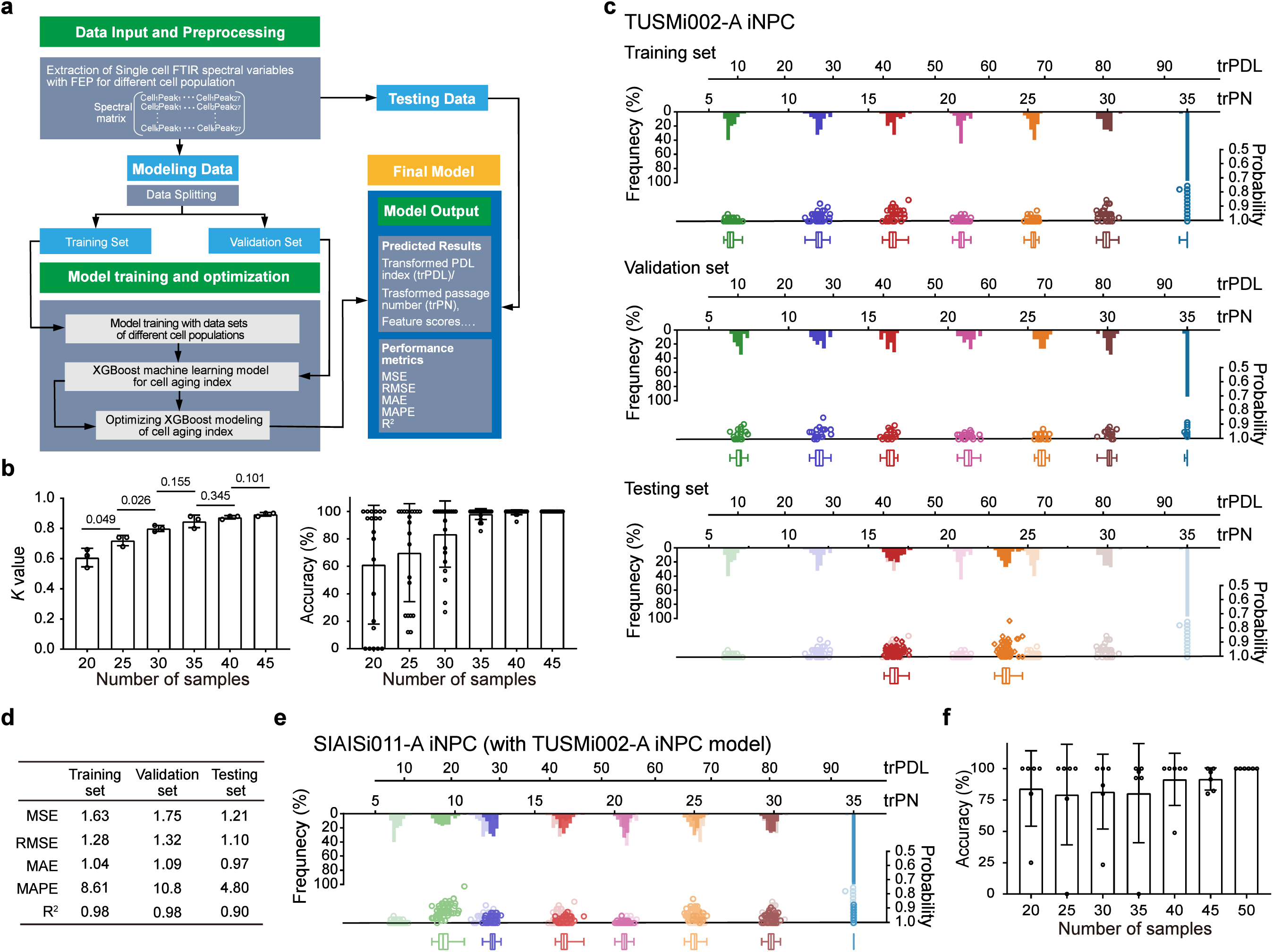
XGBoost modelling of FTIR spectral fingerprints provides trPDL/trPN as a quantitative aging index. **a.** CIADEX framework of the FTIR data processing and modelling. Single-cell FTIR spectral variables were extracted with FEP for different cell populations. Then, the data were split into training and validation sets. Models were generated and optimized with these data, utilising the XGBoost algorithm. The model was finalized with further validation using independent testing data sets. trPDL/trPN were outputted for comparison with the recorded cPDL/passage number. F scores of features, as well as performance metrics for the robustness of the model, were monitored. **b.** Model performance was assessed by comparing the predicted trPDL/trPN versus recorded cPDL/passage number. Cohen’s *K* value and the accuracy were plotted against the sample size per session. **c.** Detailed results for TUSMi002-A iNPC across three datasets (training, validation, and testing sets) were shown with raincloud plots. Dots represent the trPDL/trPN. Each set shows the distribution frequency of individual trPDL/trPN along with the corresponding probability for each given cell population. The middle line in the box plots shows the median trPDL/trPN, the hinges indicate the 25th and 75th quantiles, and the whiskers point to the 5th and 95th quantiles. **d.** Performance metrics (MSE, RMSE, MAE, MAPE, R²) across the three datasets were compared and shown. **e-f**. Generalization capabilities of the generated model. The raincloud plot represents the performance of the TUSMi002-A iNPC model for SIAISi011-A iNPCs FTIR data (e). The accuracy against sample size was shown (f).

We first evaluated the influence of training sample numbers on model stability. For TUSMi002-A iNPCs, we found that when the sample number in the training set reached 30 or more, the trPDLs aligned well with the recorded cPDLs (Figure 3b, left). Correspondingly, the dispersion of results decreased significantly, and the accuracy stabilized above 80% with 35 or more samples (Figure 3b, right). Therefore, we used 40 cells per group for the training set. The resulting model (Figure 3c) achieved an overall accuracy of 92% for the training set and 89% for the validation set (The median values and 95% confidence intervals for each group are listed in Supplementary Table 1). To test the robustness of the established model, we collected FTIR spectra from two additional culture batches of TUSMi002-A iNPCs as the independent testing data sets. The model calculated these two batches of TUSMi002-A iNPCs with an overall accuracy greater than 90% (Figure 3c and Supplementary Table 1). This model was then assessed comprehensively with Mean Squared Error (MSE), Root Mean Squared Error (RMSE), and *R*^2^. As shown in Figure 3d, these results demonstrated that the model achieved remarkable predictive performance and could provide trPDL/trPN values with *R*^2^ > 0.95 without overfit. The model for SIAISi011-A iNPCs was also established and evaluated with the same strategy (Figure S5a-c). Similar to the TUSMi002-A model, the SIAISi011-A model demonstrated high accuracy and consistent performance when the training set contained 35 or more samples (Figure S5a). The developed model performed well with training, validation, and testing datasets (Figure S5b-c and Supplementary Table 1). The relative importance of each feature in the developed models for both TUSMi002-A iNPCs and SIAISi011-A iNPCs is shown in F-score plots (Figure S5d-e). Interestingly, molecular features include C4-C5 and C=N stretching in the imidazole ring of DNA, Amide II vibrations, lipid stretching and vibrations were identified as critical discriminators for both models, indicating that structural changes in lipids, proteins, and DNA during culture cycles have been captured. Further investigation to elucidate the direct link of these spectra with membrane rigidity, proteostasis alteration, and epigenetic drift during cell aging processes should provide mechanistic insights and uncover key contributors to cellular senescence. These results indicate that the developed FTIR spectra model could provide a quantitative assessment of the aging status of given cell populations with trPDL/trPN values that were well consistent with the recorded cPDLs and passage numbers.

We then further investigated the normalization capability of the trPDL/trPN algorithm by testing whether we can apply the established models for direct side-by-side comparison of the relative aging status of different iNPC lines. To test this, SIAISi011-A iNPCs FTIR variables were subjected to trPDL/trPN value calculation using the TUSMi002-A iNPC model. Interestingly, as shown in Figure 3e, the trPDLs of SIAISi011-A iNPCs middle and later populations aligned well with the recorded cPDLs, whereas the trPDLs of early populations were higher than their recorded cPDLs, which is correlated well with the different proliferation and differentiation potency of these two cell lines. These findings indicate that, beyond clear discrimination between early and late passage cell populations using FTIR spectral signatures, differences in aging progression of individual iNPC lines from different donors could be captured and compared within this FTIR-based trPDL/trPN quantitative analysis.

Together, these results demonstrate that a cell identity and aging index (CIADEX) model could be established accordingly for individual iNPC lines utilizing the XGBoost algorithm. The trPDL/trPN generated by the CIADEX model could be applied as a quantitative aging index of individual cell populations, providing FTIR spectral features for effectively tracking the iNPC aging progress during multiple culture cycles.

### Quantification of drug-induced aging and rejuvenation of iNPCs by trPDL/trPN

Then we asked whether we could precisely capture subtle cellular aging-related phenotypes of different iNPC populations using trPDL/trPN as a quantitative aging index. To assess this potentiality, we induced aging-related phenotypes through modulation of cellular NAD metabolism, as previous studies have established that inhibition of NAD+ biosynthesis leads to premature senescence in NPCs^42^, while increased NAD+ levels promote NPC proliferation and neurogenesis^43^.

NAM and FK866 function through the provision of NAD precursor substances^40^ and the inhibition of NAMPTase^41^, respectively. TUSMi002-A iNPCs at cPLD37 (P15) were treated with various concentrations of FK866 (0.1∼3 nM) and NAM (0.1∼1 mM) for 96 hours, after which cell proliferation and NAD levels were assessed (Figure 4a and b). Treatment with 0.3 nM FK866 did not significantly affect cell proliferation while still partially reduced intracellular NAD levels, whereas 0.1 nM FK866 had no significant effect on either parameter. Conversely, both 0.3 mM and 0.1 mM NAM significantly increased intracellular NAD levels without affecting the cell proliferation rate. Based on these findings, we administered 0.3 nM FK866 or 0.3 mM NAM to the iNPC culture since cPDL37(P15) for multiple culture cycles to assess the cumulative effects of long-term treatments. As shown in Figure 4c, prolonged treatment with these micro-doses altered cell cPDL levels with the same passage numbers. Consistently, the proliferation potency of treated cells was also changed gradually, with effects becoming evident after 5 passages (i.e., 5 culture cycles, P20) and more pronounced after 10 passages (i.e., P25, Figure 4d and Figure S6a). The stemness and differentiation potential of micro-dose-treated iNPCs were also analyzed. As shown in Figure 4e and f, the expression of both Nestin and SOX2 was altered in response to both FK866 and NAM micro-dose treatments. Consistently, these compounds modulated the differentiation potential of iNPCs, affecting both the differentiation rate and the complexity of differentiated neurons, which were either reduced by FK866 or enhanced by NAM, respectively (Figure 4g-i). Similar alterations were observed in SIAISi011-A iNPCs treated with micro-doses of either FK866 or NAM (Figure S6b-h). Thus, with the prolonged micro-dose treatment of both FK866 and NAM, we observed clear aberrancy of recorded cPDLs from their correlation with culture cycles and the cellular pluripotency.

**Figure 4.**
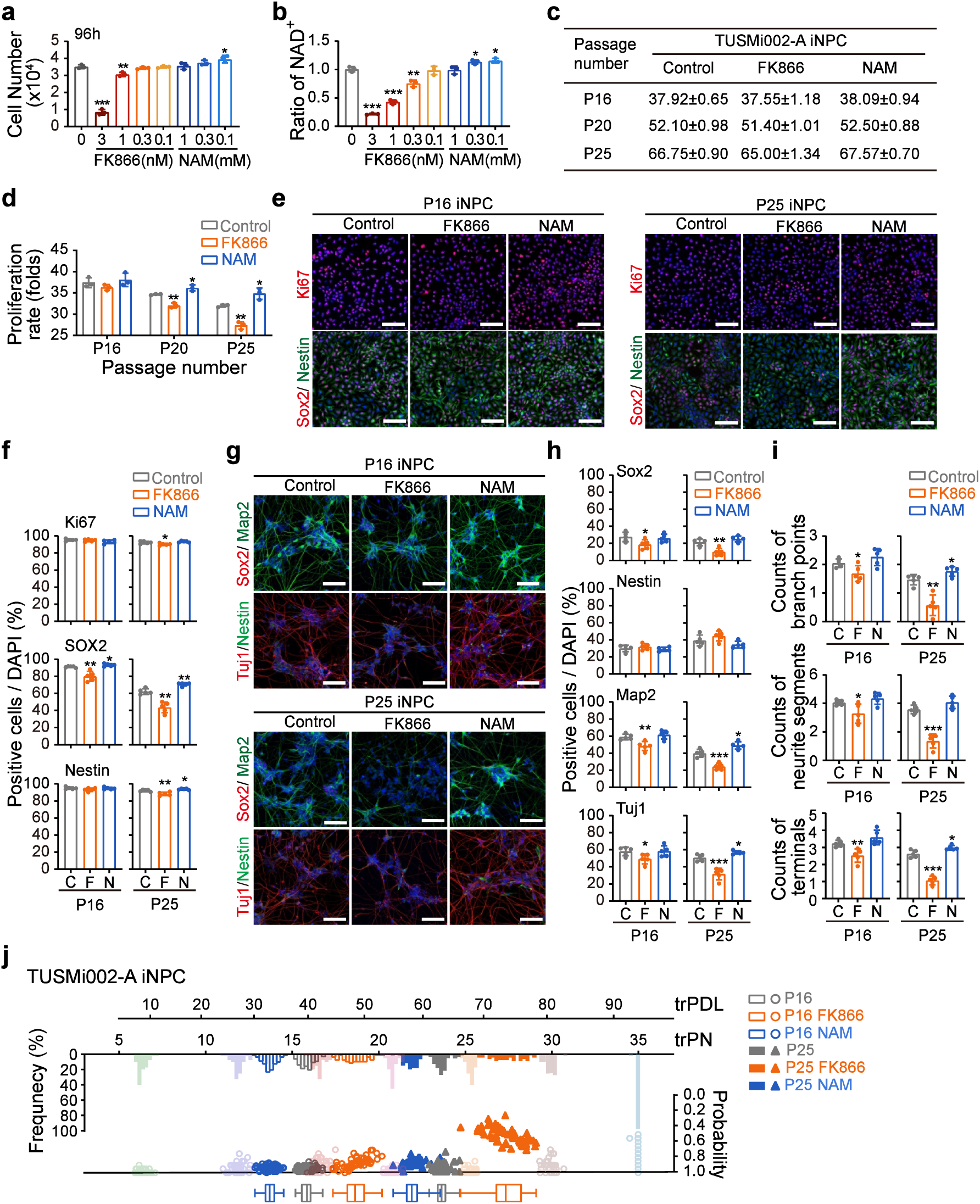
Quantitative assessment of drug-induced aging and rejuvenation phenotypes by CIADEX model. **a-b.** Modulation of iNPC proliferation and cellular NAD+ levels by FK866 or NAM treatment. TUSMi002-A iNPCs cPDL37/P15 were treated with different concentrations of FK866 or NAM as indicated. The proliferation (a) and the cellular NAD+ levels (b) in these cell populations were quantified in parallel and shown. **c**. Prolonged micro-dose treatment of FK866 or NAM altered iNPC cPDLs compared with those of the control iNPCs with the same passage number. TUSMi002-A iNPCs were treated with either 0.3 nM FK866 or 0.3 mM NAM continuously. The population expansion over multiple cultural cycles was recorded and shown. **d.** Prolonged treatment of micro-dose FK866 or NAM altered iNPC proliferation. TUSMi002-A iNPCs were treated with either 0.3nM FK866 or 0.3mM NAM for 1, 5, 10 culture cycles, and the proliferation rate was quantified and shown. **e-f**. Immunofluorescent images of Ki67, Sox2, and Nestin in TUSMi002-A iNPCs treated with either 0.3nM FK866 or 0.3mM NAM for 1 or 10 culture cycles (P16 and P25) (**e**). The percentages of Ki67, Sox2, and Nestin-positive cells were shown in parallel (**f**). Scale bar, 100 µm. **g-i**. The differentiation potency of TUSMi002-A iNPCs treated with either 0.3nM FK866 or 0.3mM NAM for 1 or 10 culture cycles (P16 and P25). Cells were induced to differentiate into neuronal cells. The morphological and immunological characteristics of neuronal cells derived from iNPCs were evaluated. Map2-positive, Tuj1-positive, Nestin-positive, or Sox2-positive cells in each group were counted **(h)**. Neuron morphology was assessed by quantitative analysis of confocal immunofluorescence microscopy images **(i)**. Specifically, the average counts of branches per neuron, neurite segments, and terminals per neuron were compared and shown. Scale bar, 100 µm. **j.** trPDL/trPN of iNPCs treated with either 0.3nM FK866 or 0.3mM NAM for 1 or 10 culture cycles (P16 and P25) were generated by the TUSMi002-A iNPC model and shown on a raincloud plot. These results are representative of at least 3 independent experiments. The data are presented as the means ± SEMs. \**P* <0.05, \*\**P* <0.01, and \*\*\**P* <0.001 compared with the control group. *P* values were determined via a two-tailed T-test.

We then obtained infrared spectral datasets of iNPCs treated with either FK866 or NAM for a single culture cycle (passage 15 to 16, normal treatment timeframe) and for 10 culture cycles (passage 15 to 25, prolonged treatment timeframe). LDA results revealed that after drug treatment, the FTIR spectral features of either NAM-treated or FK866-treated iNPCs differed from those of controls (Figure S7a-d). The trPDL/trPN values were then calculated using the established CIADEX models. As shown in Figure 4j, TUSMi002-A iNPCs treated with micro-dose FK866, which showing retarded cell proliferation (with lower recorded cPDLs compared to the control group, Figure 4c), obtained higher trPDL/trPN than the control group: trPDL46/trPN18 vs. trPDL40/trPN16 for one culture cycle treatment, and trPDL73/trPN27 vs. trPDL64/P24 for 10 culture cycle treatment. In contrast, micro-dose NAM treatment led to enhanced iNPC proliferation and increased population doubling rates with increased recorded cPDLs vs. the control group (Figure 4c). These NAM-treated iNPCs exhibited lower trPDL/trPN values compared to control iNPCs: trPDL34/trPN14 vs. trPDL40/trP16 for one culture cycle treatment, and trPDL58/P22 vs. trPDL64/P24 for 10 culture cycle treatments. Furthermore, similar results were reproduced with SIAISi011-A iNPCs treated with micro-doses of either FK866 or NAM (Figure S7e), using the CIADEX model developed for TUSMi002-A iNPCs. FK866-treatment results in accelerated trPDL/trPN: trPDL52/trPN20 versus trPDL46 /trPN18 for one culture cycle, trPDL73/P27 versus trPDL67/trPN25 for 10 culture cycles. Following NAM treatment, the iNPCs showed decelerated trPDL/trPN: trPDL37/trPN15 versus trPDL46/trPN18 for one culture cycle, and trPDL58/trPN22 versus trPDL67/trPN25 for 10 culture cycles. These findings are particularly crucial as they demonstrate that even under subtle promoting or unfavourable conditions, where recorded cPDLs and passage numbers have already aberrated and lost their correlation with cell phenotypic and functional characteristics, the trPDL/trPN values generated by the established CIADEX model remain consistent with the aging status of the cell populations.

Together, these results confirm that the CIADEX analysis framework could serve as a reliable methodology for precise comparison of cell populations under even slightly different culture conditions. Furthermore, given the ability of the CIADEX analysis framework to effectively track cell aging progress across different cell populations under various culture conditions, trPDL/trPN values could be valuable evaluation metrics for cell culture optimization as well as related drug screening.

## Discussion

While prior FTIR microspectroscopy offers molecular fingerprints of cells but lacks standardized analytical pipelines, our results show that CIAdex can leverage machine learning to decode these spectral fingerprints for characterizing cell populations and quantifying aging. We demonstrated, for the first time, that FTIR spectral fingerprints, which decipher structural biomolecular shifts as well as compositional changes, can be applied to differentiate biological variations across multiple levels - from cell types, cell lines, to subtle batch and passage differences. Crucially, trPDL/trPN, its aging index, calculated from FTIR data via the XGBoost algorithm, reliably tracks aging dynamics, responding to both natural and drug-induced cellular senescence or rejuvenation, outperforming conventional metrics like cPDLs or static senescence marker panels. Thus, CIAdex resolves two fundamental challenges in cell therapy development: rapid identity validation and real-time aging monitoring.

Notably, longitudinal tracking of iNPCs over 35 passages confirmed robustness of the trPDL/trPN, aligning with aging hallmarks such as membrane rigidity, epigenetic shifts, proteostasis alteration, and metabolic decline. Furthermore, the sensitivity of trPDL/trPN to subtle drug-modulated aging (e.g., NAD+ regulators) highlights its potential to act as universal aging indicators, which could further deepen our insights into cellular senescence and rejuvenation, and thereby support the development of anti-aging strategies. Thus, by transforming infrared spectroscopy from a compositional analysis tool into a dynamic label-free cellular phenotyping framework, CIAdex supports aging research by quantifying biological age through spectral markers, expanding the multidimensional cellular aging atlases, and complementing current tools such as epigenetic clocks by Hartmann et al^19^.

Cell therapy manufacturing requires balancing sufficient cell expansion for effective dosing with the prevention of replicative senescence that impairs function. CIAdex tackles this by offering in-process quality monitoring, replacing otherwise labor-intensive, time-consuming, and low-throughput assays. By capturing distinct molecular fingerprints through FTIR microspectroscopy, CIAdex enables precise label-free discrimination of diverse manufacturing batches and culture cycles - a critical capability for rigorous standardization of cell therapy products. Furthermore, the trPDL/trPN index provides a quantitative aging metric to establish quality benchmarks. While our current models are generated and validated for iNPCs, CIAdex’s adaptability extends to other cell types such as mesenchymal stromal cells and hematopoietic stem cells. Thus, an open-access FTIR spectral database could amplify CIAdex’s utility and support its future clinical translation. Moreover, our work also suggests possibilities for translating laboratory-based spectral analysis into field-deployable devices in the future, directly complying with emerging stringent regulatory requirements on in-process monitoring of cell-based product manufacturing.

## Materials and Methods

### Cell lines and cell culture

The human neural progenitor cells used in this study were generated from TUSMi002-A^29^, TUSMi006-A^35^, TUSMi007-A^36^, and SIAISi011-A^37^ human induced pluripotent stem cells (iPSCs) derived from healthy volunteers, which have been registered in hPSCreg (https://hpscreg.eu) with documents of ethical approval and informed consent.

Human iPSCs were cultured and passaged in mTeSR™ on Matrigel Matrix (354277, Corning Inc., Bedford, MA, USA) or poly-L-ornithine and laminin precoated plates. hNPCs were induced from iPSCs using a neural induction medium according to the monolayer culture protocols suggested by the manufacturer (STEMCELL Technologies, Columbia, Canada). After 10 days of induction in the neural induction medium, the cells were transferred to the neural progenitor medium. The generated hNPCs were further cultured to confluence and then expanded and passaged. Alternatively, hNPCs were cultured in NPC basal medium containing DMEM/F-12, neurobasal medium, N2 supplement, B27 supplement minus vitamin A, NEAA, or GlutaMAX. CHIR99021 (3 μM), SB431542 (5 μM), and bFGF (10 ng/ml) were used as reported previously^58–60^. For differentiation of hNPCs, 30,000 cells per well were seeded on 10 µg/ml poly-L-ornithine (PLO) and 10 µg/ml laminin-coated 24-well plates in neural progenitor medium. Two hours later, the medium was changed to neuron differentiation medium containing neurobasal medium, N2 supplement, B27 supplement, culture media, 0.1 μM cAMP, 200 µg/ml L-AA, 10 ng/ml BDNF, 10 ng/ml GDNF, and 10 ng/ml IGF-1. The medium was changed every 2-3 days. All the cells were cultured at 37□ with 5% CO_2_ in a humidified atmosphere.

Cell viability was consistently 94%-98 %, as determined by Trypan blue exclusion using an Automated Image-Based Cell Analyzer (Countstar Mira BF, Shanghai Ruiyu Biotech Ltd., China). The population doubling level (PDL) per each culture cycle (i.e., population doubling rate) was calculated as: PDL*_i_* = log_2_(*N_ei_*/*N_bi_*) = 3.32(log*N_ei_*-log*N_bi_*), where *N_ei_* is the final viable cell count, and *N_bi_*is the initial seeded cell count of passage number *i* (the *i-*th culture cycle). cPDL (cumulative population doubling level) was calculated as: cPDL*_i_* = PDL*_i_* + PDL_(*i-1)*_ = 3.32(log*N_ei_*– log*N_0_*) + *S*, where *N_0_* is the initial seeded cell count and *S* is the recorded PDL at the start of the culture procedure^61^.

### Chemicals, antibodies, culture media, and kits

Poly-L-ornithine solution (PLO, P4957), Laminin (L2020), L-Ascorbic Acid (L-AA, Vitamin C, A4403), and DAPI (D9542) were obtained from Sigma Aldrich (St. Louis, MO, USA). Daporinad (FK866, HY-50876) and Nicotinamide (NAM, HY-B0150) were obtained from MedChemExpress (Monmouth Junction, NJ, USA).

The antibodies used in this study include: Oct3/4 Antibody (C-10) (sc5279, Santa Cruz Biotechnology, CA, USA), Nanog (D73G4) XP Rabbit mAb (4903S, Cell Signaling Technology, Danvers, MA, USA), SSEA-4 Antibody (813-70) (sc21704, Santa Cruz Biotechnology), Sox2 (L1D6A2) Mouse mAb (4900S, Cell Signaling Technology), Purified Mouse Anti-Nestin (61158, BD Biosciences), anti-Ki67 antibody (ab15580, Abcam), Anti-β-Tubulin III antibody produced in rabbit (Tuj1, T2200, Sigma-Aldrich), rabbit anti-Map2 (PA585755, Thermo fisher), TRA1-81 Antibody (4745S, Cell Signaling Technology, Danvers, MA, USA), TRA1-60 Antibody (ab16288, Abcam), AFP Antibody (A8452, Sigma-Aldrich), SMA Antibody (A2547, Sigma-Aldrich). Alexa Fluor 488/594/647-conjugated donkey anti-mouse and anti-rabbit IgG secondary antibodies (A21202, A21203, A21206, A21207) were obtained from Life Technologies (Eugene, OR, USA).

mTeSR™ 1 (85850), STEMdiff^TM^ Neural Induction Medium (05835), Neural Progenitor Medium (05833), Human Recombinant bFGF (78003.1), Human Recombinant BDNF Cytokines (78005.1), Human Recombinant GDNF (78058.1), and Human Recombinant IGF-1 (78022.1) were obtained from STEMCELL Technologies (Columbia, Canada). Dulbecco’s modified Eagle’s medium/F-12 mixture (DMEM/F-12, 1132003), Neurobasal Medium (21103049), N2 Supplement (17502048), B-27 Supplement, minus vitamin A (12587010), B-27 Supplement, serum-free (17504044), Non-Essential Amino Acids Solution (NEAA, 11140050), GlutaMAX (35050061) and CultureOne (A3320201) were obtained from Gibco (Grand Island, NY, USA).

### Immunofluorescence staining, microscopy and high-content analysis

For in vitro immunostaining, cells cultured in multiple-well plates or coverslips were fixed with 4% PFA for 15 min at 37°C. After being washed with PBS three times, the samples were transferred to a blocking buffer (1% bovine serum albumin and 0.5% Triton X-100 in PBS) and incubated at room temperature for 30 min. Then, the samples were incubated with primary antibodies at 4°C overnight. The dilution ratios of the following primary antibodies were optimized for signal-to-noise ratios: Oct3/4 (1:500), Nanog (1:500), SSEA4 (1:500), Sox2 (1:200), Nestin (1:500), Ki67 (1:200), Tuj1 (1:200), Map2 (1:200), TRA1-81 (1:200), TRA1-60 (1:200), AFP (1:200), and SMA (1:100). After being washed 3 times, the samples were further incubated with the corresponding secondary antibodies (Alexa Fluor 488/561/647-conjugated donkey anti-mouse or anti-rabbit IgG, 1:1000) at room temperature for two hours. DAPI was used for counterstaining the nuclei. A confocal laser scanning microscope (Carl Zeiss LSM710, Jena, Germany) was used for cell image capture. Alternatively, images were acquired on an Operetta CLS ™ High-Content Analysis microscope imaging System (Perkin Elmer Operetta CLS, Waltham, USA) using a 20× objective lens. Quantitative image analysis was performed using Harmony 3.5.2 software (Perkin Elmer, Waltham, USA).

### NAD measurement

The nicotinamide adenine dinucleotide (NAD) was measured using the commercial NAD/NADH Quantification Kit (Sigma-Aldrich MAK037). In brief, cells were collected and then lysed in NADH/NAD extraction buffer. After the proteins were deactivated and removed, the supernatants were mixed with the Master Reaction Mix and incubated at room temperature for 5 minutes in a 96-well plate. Then, the color developer for NADH was added. When the color had been properly developed, the reaction was terminated by adding the stop solution. The absorbance at 450 nm was recorded. The relative NAD content was calculated for the drug-treated and untreated groups according to the standard curve.

### Synchrotron Radiation FTIR Microspectroscopy Experiments and Data Analysis

Cells were seeded onto CaF_2_ chips pre-coated with Matrigel. Cells were fixed with 4% paraformaldehyde, washed with ultrapure water, and completely dried at room temperature. SR-FTIR was carried out at the beamline BL01B of Shanghai Synchrotron Radiation Facility (SSRF), which is equipped with a Nicolet 6700 Fourier transform infrared spectrometer, a Nicolet Continuμm infrared microscope with a liquid N_2_ cooled MCT detector, and a 32× infrared objective^62^. To ensure the collected spectra of single cells with comparable signal intensities and signal/noise ratio, the same aperture size of 25 × 25 μm was used throughout the experiment. All the spectra were obtained within the mid-infrared region of 4000-1000 cm^−1^ with a resolution of 4 cm^−1^ and 64 coadded scans for each spectrum. Original spectra were smoothed (15-point) and corrected with a linear automatic baseline. The second-derivative spectra were calculated using the Savitzky-Golay algorithm. All data collection and pre-processing procedures were carried out using OMNIC (Thermo Fisher Scientific Inc.). Infrared spectral Mie scattering was corrected on MATLAB R2022a using Resonant Mie Scattering EMSC (RMieS-EMSC) correction. Linear Discriminant Analysis (LDA) was also carried out on MATLAB and plotted using Origin 2024.

For collecting single-cell infrared mapping images, the step size was set to 2 μm, and there were 32 coadded scans for each spectrum. After correcting for Mie Scatter, the mapping images were smoothed (15-point), baseline corrected (linear method), and normalized (offset) on CytoSpec. Fatty acid, protein, and nucleic acid components were used to construct the chemical image of the cells based on the areas of 3000-2830^−1^, 1770-1475^−1^, and 1300-1000 cm^−1^, respectively.

### The Cell Identity and Aging Index (CIADEX) Framework

Cellular feature preprocessing and model construction were performed through the following steps: 1. Data Preprocessing and Feature Extraction: Key cellular feature peaks were extracted from the modelling cohort using random sampling. Statistical descriptors, including mean, median, and variance, were calculated for each characteristic peak to establish comprehensive feature profiles. 2. Model Construction and Training: The XGBoost regression model is implemented using the XGBoost package (Python, Anaconda3, Jupyter notebook). The model architecture employed 100 decision trees with maximum depth constrained to 6 layers, optimized through 5-fold cross-validation. Training utilized adaptive learning rates (η=0.01-0.3) with L2 regularization (λ=1.0) to prevent overfitting. 3. Single-cell Generation Prediction: For validation cohort analysis, each target cell was randomly combined with 9 reference cells to form cell groups (n=100 permutations per target). The trained model generated probabilistic predictions for each permutation, with the final generation assignment determined by the median prediction value across all permutations. Generation probabilities were estimated by integrating the area under the prediction density curve within ±2 generation ranges from the median value.

## QUANTIFICATION AND STATISTICAL ANALYSIS

Statistical analysis was performed using GraphPad Prism 7. Unless stated otherwise, all the data are presented as the mean ± standard error of the mean (SEM). The value of n for sample size and replicates can be found in the figure legends. All the statistical tests used are also indicated in the respective figure legends. A t-test was used to determine the statistical significance of differences between the two groups (* for *P* < 0.05; ** for *P* < 0.01, and *** for *P* < 0.001).

## Data Availability Statement

### Lead contact

Further information and requests for resources and reagents should be directed to and will be fulfilled by the lead contact, Ying Wang.

### Materials Availability

All unique biological materials generated in this study are available upon request with a completed Materials Transfer Agreement.

### Data and Code Availability

All RNA sequencing data of this study have been deposited in the National Genomics Data Center under accession numbers PRJCA036864. Codes generated in this study have been deposited at https://github.com/Neuralrepair/CIADEX. Any additional information required to reanalyze the data reported in this paper is available from the lead contact upon request.

## Acknowledgements

We thank all members of the laboratory for sharing reagents and ideas. We thank the members of the core facility of ShanghaiTech University for their technical assistance. We thank the staff members of the BL01B beamline (https://cstr.cn/31129.02.NFPS.BL01B) at the National Facility for Protein Science in Shanghai (https://cstr.cn/31129.02.NFPS) for providing technical support and assistance in data collection and analysis. This work was supported, in part, by grants 2022YFA1105003 and 2023YFC2506100 from the National Key Research and Development Program of China, grant YDZX20213100001684 from the Shanghai Science and Technology Development Foundation, and grant 82171400 from the National Natural Science Foundation of China.

## Author Contributions

JZ: supervision, conception and design, data analysis and interpretation, manuscript writing, and final approval of the manuscript; YW, YD and CH: performed the experiments, collection of the data, data analysis and interpretation, and manuscript preparation; XZ: data collection and analysis; JT and YD: contribution of analysis systems, final approval of the manuscript.

## References

1. Aijaz, A. et al. Biomanufacturing for clinically advanced cell therapies. Nat Biomed Eng 2, 362–376 (2018).

2. Zhang, Z. et al. A panoramic view of cell population dynamics in mammalian aging. Science 387, eadn3949 (2025).

3. Kirkeby, A., Main, H. & Carpenter, M. Pluripotent stem-cell-derived therapies in clinical trial: A 2025 update. Cell Stem Cell 32, 10–37 (2025).

4. Wagoner, Z. W. & Zhao, W. Therapeutic implications of transplanted-cell death. *Nat*. Biomed. Eng. 5, 379–384 (2021).

5. Bonder, M. J. et al. scEpiAge: an age predictor highlighting single-cell ageing heterogeneity in mouse blood. Nat. Commun. 15, 7567 (2024).

6. Schiff, L. et al. Integrating deep learning and unbiased automated high-content screening to identify complex disease signatures in human fibroblasts. Nat. Commun. 13, 1590 (2022).

7. Yu, D. et al. CellBiAge: Improved single-cell age classification using data binarization. Cell Rep. 42, 113500 (2023).

8. Yin, J. Q., Zhu, J. & Ankrum, J. A. Manufacturing of primed mesenchymal stromal cells for therapy. Nat Biomed Eng 3, 90–104 (2019).

9. Harley, J. et al. Telomere shortening induces aging-associated phenotypes in hiPSC-derived neurons and astrocytes. Biogerontology 25, 341–360 (2024).

10. Duscher, D. et al. Aging disrupts cell subpopulation dynamics and diminishes the function of mesenchymal stem cells. Sci Rep 21, 07144 (2014).

11. Andrew, T., David, E. & Raymond, W. N. Q5D Derivation and Characterization of Cell Substrates Used for Production of Biotechnological/Biological Products. ICH Qual. Guidel. Ch13, (2017).

12. Geraghty, R. J. et al. Guidelines for the use of cell lines in biomedical research. Br. J. Cancer 111, 1021–1046 (2014).

13. Food and Drug Administration. Regulatory Considerations for Human Cells, Tissues, and Cellular and Tissue-Based Products: Minimal Manipulation and Homologous Use; G. *Uidance Ind*. Food Drug Adm. Staff (2017).

14. Liu, Y. et al. Mapping Cell Phenomics with Multiparametric Flow Cytometry Assays. Phenomics 2, 272–281 (2022).

15. Lazarski, C. A. & Hanley, P. J. Review of flow cytometry as a tool for cell and gene therapy. Cytotherapy 26, 103–112 (2024).

16. Vandereyken, K., Sifrim, A., Thienpont, B. & Voet, T. Methods and applications for single-cell and spatial multi-omics. Nat Rev Genet 24, 494–515 (2023).

17. Gulati, G. S., D’Silva, J. P., Liu, Y., Wang, L. & Newman, A. M. Profiling cell identity and tissue architecture with single-cell and spatial transcriptomics. Nat Rev Mol Cell Biol 26, 11–31 (2025).

18. David Bernard, Emmanuel Doumard, Isabelle Ader, Philippe Kemoun, Jean-Christophe Pagès, Anne Galinier, Sylvain Cussat-Blanc, Felix Furger, Luigi Ferrucci, Julien Aligon, Cyrille Delpierre, Luc Pénicaud, Paul Monsarrat, Louis Casteilla. Explainable machine learning framework to predict personalized physiological aging. Aging Cell 22, 13872 (2023).

19. Hartmann, C. et al. Systematic estimation of biological age of in vitro cell culture systems by an age-associated marker panel. Front Aging 4, 1129107 (2023).

20. Leslie, M. Infrared method could safely identify cells. Science 361, 541 (2018).

21. Marcelli, A., Cricenti, A., Kwiatek, W. M. & Petibois, C. Biological applications of synchrotron radiation infrared spectromicroscopy. Biotechnol Adv 30, 1390–404 (2012).

22. Hughes, C. et al. SR-FTIR spectroscopy of renal epithelial carcinoma side population cells displaying stem cell-like characteristics. Analyst 135, 3133–41 (2010).

23. Nakamura, T. et al. Microspectroscopy of spectral biomarkers associated with human corneal stem cells. Mol Vis 16, 359–68 (2010).

24. Theophilou, G. et al. Synchrotron- and focal plane array-based Fourier-transform infrared spectroscopy differentiates the basalis and functionalis epithelial endometrial regions and identifies putative stem cell regions of human endometrial glands. Anal Bioanal Chem 410, 4541–4554 (2018).

25. Liu, Z. et al. Synchrotron FTIR microspectroscopy reveals early adipogenic differentiation of human mesenchymal stem cells at single-cell level. Biochem Biophys Res Commun 478, 1286–91 (2016).

26. Chonanant, C. et al. Characterisation of chondrogenic differentiation of human mesenchymal stem cells using synchrotron FTIR microspectroscopy. Analyst 136, 2542–51 (2011).

27. Tiwari, S. et al. INFORM: INFrared-based ORganizational Measurements of tumor and its microenvironment to predict patient survival. Sci Adv 7, (2021).

28. Miller, L. M. & Dumas, P. Chemical imaging of biological tissue with synchrotron infrared light. Biochim Biophys Acta 1758, 846–57 (2006).

29. Wang, Y. et al. Generation of a human induced pluripotent stem cell line from a 65-year old healthy female donor with Chinese Han genetic background. Stem Cell Res 24, 33–35 (2017).

30. Wang, Y. et al. Generation of a human control PBMC derived iPS cell line TUSMi001-A from a healthy male donor of Han Chinese genetic background. Stem Cell Res 25, 22–25 (2017).

31. Aksoy, C. & Severcan, F. Role of Vibrational Spectroscopy in Stem Cell Research. *Spectrosc*. Int. J. 27, 167–184 (2012).

32. Aksoy, C., Guliyev, A., Kilic, E., Uckan, D. & Severcan, F. Bone marrow mesenchymal stem cells in patients with beta thalassemia major: molecular analysis with attenuated total reflection-Fourier transform infrared spectroscopy study as a novel method. Stem Cells Dev 21, 2000–11 (2012).

33. Jackson, M. & Mantsch, H. H. Medical Science Applications of IR. Encycl. Spectrosc. Spectrom. Second Ed. 1494–1502 (1999) doi:10.1016/B978-0-12-374413-5.00200-1.

34. Li, Z., Nie, F., Wu, D., Wang, Z. & Li, X. Sparse Trace Ratio LDA for Supervised Feature Selection. IEEE Trans Cybern 54, 2420–2433 (2024).

35. Wang, Y. et al. Derivation of induced pluripotent stem cells TUSMi006 from an 87-year old Chinese Han Alzheimer’s disease patient carrying GRINB and SORL1 mutations. Stem Cell Res 31, 127–130 (2018).

36. Wang, Y., Zhang, J., Lei, Y. & Zhao, J. Establishment of TUSMi007-A, an induced pluripotent stem cell (iPSC) line from an 83-year old Chinese Han patient with Alzheimer’s disease (AD). Stem Cell Res 33, 265–268 (2018).

37. Zhang, W. et al. Generation of a human induced pluripotent stem cell line (SIAISi011-A) from a 61-year-old Chinese Han healthy female donor. Stem Cell Res 51, 102173 (2021).

38. Krüger, A., Bürkle, A., Hauser, K. & Mangerich, A. Real-time monitoring of PARP1-dependent PARylation by ATR-FTIR spectroscopy. Nat Commun 11, 2174 (2020).

39. Wiese, D. M., Ruttan, C. C., Wood, C. A., Ford, B. N. & Braid, L. R. Accumulating Transcriptome Drift Precedes Cell Aging in Human Umbilical Cord-Derived Mesenchymal Stromal Cells Serially Cultured to Replicative Senescence. Stem Cells Transl Med 8, 945–958 (2019).

40. Liu, D. et al. Nicotinamide forestalls pathology and cognitive decline in Alzheimer mice: evidence for improved neuronal bioenergetics and autophagy procession. Neurobiol Aging 34, 1564–80 (2013).

41. Gerner, R. R. et al. NAD metabolism fuels human and mouse intestinal inflammation. Gut 67, 1813–1823 (2018).

42. Stein, L. R. & Imai, S. Specific ablation of Nampt in adult neural stem cells recapitulates their functional defects during aging. Embo J 33, 1321–40 (2014).

43. Hou, Y., et al. NAD(+) supplementation normalizes key Alzheimer’s features and DNA damage responses in a new AD mouse model with introduced DNA repair deficiency. Proc Natl Acad Sci U A 115, E1876–e1885 (2018).

44. Mitch, L. Researchers coax stripped-down cells to grow normally. Science 372, 18 (2021).

45. Oh, J., Lee, Y. D. & Wagers, A. J. Stem cell aging: mechanisms, regulators and therapeutic opportunities. Nat Med 20, 870–80 (2014).

46. Levine, M. E. et al. An epigenetic biomarker of aging for lifespan and healthspan. Aging 10, 573–591 (2018).

47. Blasco, M. A. Telomere length, stem cells and aging. Nat Chem Biol 3, 640–9 (2007).

48. Moqri, M. et al. Biomarkers of aging for the identification and evaluation of longevity interventions. Cell 186, 3758–3775 (2023).

49. Rossiello, F., Jurk, D., Passos, J. F. & d’Adda di Fagagna, F. Telomere dysfunction in ageing and age-related diseases. Nat. Cell Biol. 24, 135–147 (2022).

50. Brunet, A., Goodell, M. A. & Rando, T. A. Ageing and rejuvenation of tissue stem cells and their niches. Nat Rev Mol Cell Biol 24, 45–62 (2023).

51. Bernadotte, A., Mikhelson, V. M. & Spivak, I. M. Markers of cellular senescence. Telomere shortening as a marker of cellular senescence. Aging 8, 3–11 (2016).

52. Argentieri, M. A. et al. Proteomic aging clock predicts mortality and risk of common age-related diseases in diverse populations. Nat. Med. 30, 2450–2460 (2024).

53. Schneider, E. L. & Mitsui, Y. The relationship between in vitro cellular aging and in vivo human age. Proc. Natl. Acad. Sci. 73, 3584–3588 (1976).

54. Kuo, C.-L. et al. Proteomic aging clock (PAC) predicts age-related outcomes in middle-aged and older adults. Aging Cell 23, e14195 (2024).

55. Smith, J. R., et al. Relationship Between In Vivo Age and In Vitro Aging: Assessment of 669 Cell Cultures Derived From Members of The Baltimore Longitudinal Study of Aging. J. Gerontol. A. Biol. Sci. Med. Sci. 57, B239–B246 (2002).

56. Mao, S. et al. A transcriptome-based single-cell biological age model and resource for tissue-specific aging measures. Genome Res. 33, 1381–1394 (2023).

57. Patricia R. Pitrez, Luis M. Monteiro, Oliver Borgogno, Xavier Nissan, Jerome Mertens & Lino Ferreira. Cellular reprogramming as a tool to model human aging in a dish. Nat Commun 15, 1816 (2024).

58. Brewer, G. J. & Torricelli, J. R. Isolation and culture of adult neurons and neurospheres. Nat. Protoc. 2, 1490–1498 (2007).

59. Hu, W. et al. Direct Conversion of Normal and Alzheimer’s Disease Human Fibroblasts into Neuronal Cells by Small Molecules. Cell Stem Cell 17, 204–212 (2015).

60. Cheng, L. et al. Generation of neural progenitor cells by chemical cocktails and hypoxia. Cell Res. 24, 665–679 (2014).

61. Lennon, D. P., Schluchter, M. D. & Caplan, A. I. The effect of extended first passage culture on the proliferation and differentiation of human marrow-derived mesenchymal stem cells. Stem Cells Transl. Med. 1, 279–288 (2012).

62. Wang, Y. et al. Single-Cell Infrared Microspectroscopy Quantifies Dynamic Heterogeneity of Mesenchymal Stem Cells during Adipogenic Differentiation. Anal Chem 93, 671–676 (2021).

