## supplemental figures 1-7 and supplemental table 1 for "CIAdex: Single-Cell FTIR Spectral Fingerprinting for Cell Identity Verification and Aging Quantification in Therapeutic Cell Manufacturing"

#### Supplemental Figure Legends

**Figure S1. Characterization of iPSCs and iNPCs. (related to Figure 1)**

**a, b.** Characterization of TUSMi002-A iPSCs, TUSMi006-A iPSCs, TUSMi007-A iPSCs, and SIAISi011-A iPSCs. Immunofluorescent images of stemness markers were shown (a). The pluripotency of hiPSCs was further confirmed by directed differentiation into cells of all three germ layers: endoderm (stained against alpha-fetoprotein, AFP), mesoderm (stained against smooth muscle actin, SMA), and ectoderm (stained against Nestin) (b).

**c**. Characterization of iNPCs derived from iPSCs with neural progenitor markers.

Cell nuclei were labeled with DAPI (blue). Scale bars, 100 μm.

**Figure S2. Infrared spectral absorption bands of stem cells. (related to Figure 1)**

**a.** FTIR absorption spectrum, the first and second derivatives.

**b**. Detailed information on infrared spectral absorption bands included in this study.

**Figure S3. FTIR spectral variables reveal donor- and batch-specific biochemical alterations in iNPCs. (related to Figure 1)**

3D scatter plots of LDA cluster analysis of different batches from TUSMi002-A iNPCs (labeled as TUSMi002-A-1 to TUSMi002-A-5) with SIAISi011-A iNPCs (**a**), TUSMi006-A iNPCs (**b**) or TUSMi007-A iNPCs (**c**). The left plots showed clear separations between different cell populations. The right plots show F scores for the wavenumber and height of each peak. The peak numbers for the five features with the highest F-scores were shown.

**Figure S4. FTIR spectral variables reveal biochemical alterations in iNPC with different cPDL. (related to Figure 2)**

**A.** Monitoring population expansion of SIAISi011-A iNPCs over a prolonged period in culture. Top: cPDL (cumulative population doubling level) was plotted against passage number, showing a quadratic relationship ($y = ax^2 + bx + c$) with a strong correlation ($R^2 = 0.999$). Bottom: population doubling rate (i.e., population doubling level per each culture cycle) logarithmically decreased as passage number increased, following a linear decay model ($y = dx + e$).

**b-c**. Immunofluorescent images of Ki67, Sox2, and Nestin in SIAISi011-A iNPCs across four different cPDL/passage points: cPDL4/P5, cPDL37/P15, cPDL67/P25, and cPDL93/P35 (**b**). The percentages of Ki67, Sox2, and Nestin-positive cells across the four cPDL points were shown (**c**). Scale bars, 100 μm.

**d**. The proliferation of SIAISi011-A iNPCs across the four cPDL points was shown and compared.

**e-g**. The differentiation potency of SIAISi011-A iNPCs across the four cPDL points. Cells at the indicated cPDL points were induced to differentiate into neuronal cells. The morphological and immunological characteristics of neuronal cells derived from iNPCs were evaluated. Map2-positive, Tuj1-positive, Nestin-positive, or Sox2-positive cells in each group were counted **(f)**. Neural morphology was assessed by quantitative analysis of immunofluorescent images captured via confocal microscopy; specifically, the average counts of branches per neuron, neurite segments, and terminals per neuron were compared and shown **(g)**. Scale bars, 100 μm.

**h**. The plot shows F scores for the wavenumber and height of each peak corresponding to the LDA clustering analysis of TUSMi002-A iNPC populations across different cPDL points. The peak numbers for the five features with the highest F-score were shown.

**i**. FTIR spectral data and the second derivatives for SIAISi011-A iNPC at different cPDL points were shown.

**j**. 3D scatter plots of LDA analysis showing clear clustering of SIAISi011-A iNPC populations across different cPDL points.

These results were representative of at least 3 independent experiments. Data were presented as the means ± SEMs. *$P$ <0.05, **$P$ <0.01, and ***$P$ <0.001 compared to the cPDL4 group. $P$ values were determined by a two-tailed T-test.

**Figure S5. XGBoost modelling of SIAISi011-A FTIR Spectral fingerprints. (related to Figure 3)**

**a**. Model performance was assessed by comparing the predicted trPDL/trPN versus the recorded cPDL/passage number. Cohen's $K$ value and the accuracy were plotted against the sample size per session.

**b**. Detailed results for SIAISi011-A iNPC across three datasets (training, validation, and testing sets) were shown with raincloud plots. Dots represent the trPDL/trPN. Each set shows the distribution frequency of individual trPDL/trPN along with the corresponding probability for different cell populations. The middle line in the box plots shows the median trPDL/trPN, the hinges indicate the 25th and 75th quantiles, and the whiskers point to the 5th and 95th quantiles.

**c**. Performance metrics (MSE, RMSE, MAE, MAPE, $R^2$) across the three datasets were compared and shown.

**d-e**. F scores for the wavenumber and height of each peak corresponding to the CIADEX model of TUSMi002-A iNPC (d) and SIAISi011-A iNPC (e) were sorted and shown.

**Figure S6. Drug-induced aging phenotypes (related to Figure 4)**

**a**. The proliferation curve of TUSMi002-A iNPCs treated with either 0.3nM FK866 or 0.3mM NAM for 1, 5 or 10 culture cycles (P16, P20 and P25) were monitored and shown.

**b-c.** Modulation of iNPC proliferation and cellular NAD+ levels by FK866 or NAM treatment. SIAISi011-A iNPCs cPDL37/P15 were treated with different concentrations of FK866 or NAM as indicated. The number of proliferating iNPCs was quantified and shown (b). The cellular NAD+ levels in these cell populations were monitored in parallel (c).

**d**. Prolonged treatment of micro-dose FK866 or NAM altered iNPC cPDLs compared with those of the control iNPCs with the same passage number. SIAISi011-A iNPCs were treated with either 0.3 nM FK866 or 0.3 mM NAM continuously. The population expansion over multiple cultural cycles was recorded and shown.

**e-f.** Prolonged treatment of micro-dose FK866 or NAM altered iNPC proliferation. SIAISi011-A iNPCs were treated with either 0.3nM FK866 or 0.3mM NAM for 1, 5, 10 culture cycles, and the number of proliferating iNPCs was quantified and shown.

**g**. Immunofluorescent images of Ki67, Sox2, and Nestin in SIAISi011-A iNPCs treated with either 0.3nM FK866 or 0.3mM NAM for 1 or 10 culture cycles (P16 and P25) (left). The percentages of Ki67, Sox2, and Nestin-positive cells were shown in parallel (right). Scale bar, 200 μm.

**h**. The differentiation potency of SIAISi011-A iNPCs treated with either 0.3nM FK866 or 0.3mM NAM for 1 or 10 culture cycles (P16 and P25). Cells were induced to differentiate into neuronal cells. The morphological and immunological characteristics of neuronal cells derived from iNPCs were measured. Map2-positive, Tuj1-positive, Nestin-positive, or Sox2-positive cells in each group were counted. Neural morphology was assessed by quantitative analysis of confocal immunofluorescence microscopy images. Specifically, the average number of branches per neuron, the average number of terminal orders and terminals per neuron were measured and compared. Scale bar, 100 μm.

These results are representative of at least 3 independent experiments. The data are presented as the means ± SEMs. *$P$ <0.05, **$P$ <0.01, and ***$P$ <0.001 compared with the control group. $P$ values were determined via a two-tailed T-test.

**Figure S7. Quantitative assessment of micro-dosing drug-induced iNPC aging phenotypes by CIADEX model. (related to Figure 4)**

**a-d**. 3D scatter plots of LDA analysis showing clear clustering of iNPC populations treated with either 0.3nM FK866 or 0.3mM NAM for 1 or 10 culture cycles (P16 and P25).

**e.** trPDL/trPN of iNPCs treated with either 0.3nM FK866 or 0.3mM NAM for 1 or 10 culture cycles (P16 and P25) were generated by the SIAISi011-A iNPCs model and shown on a raincloud plot.

#### Figure S1

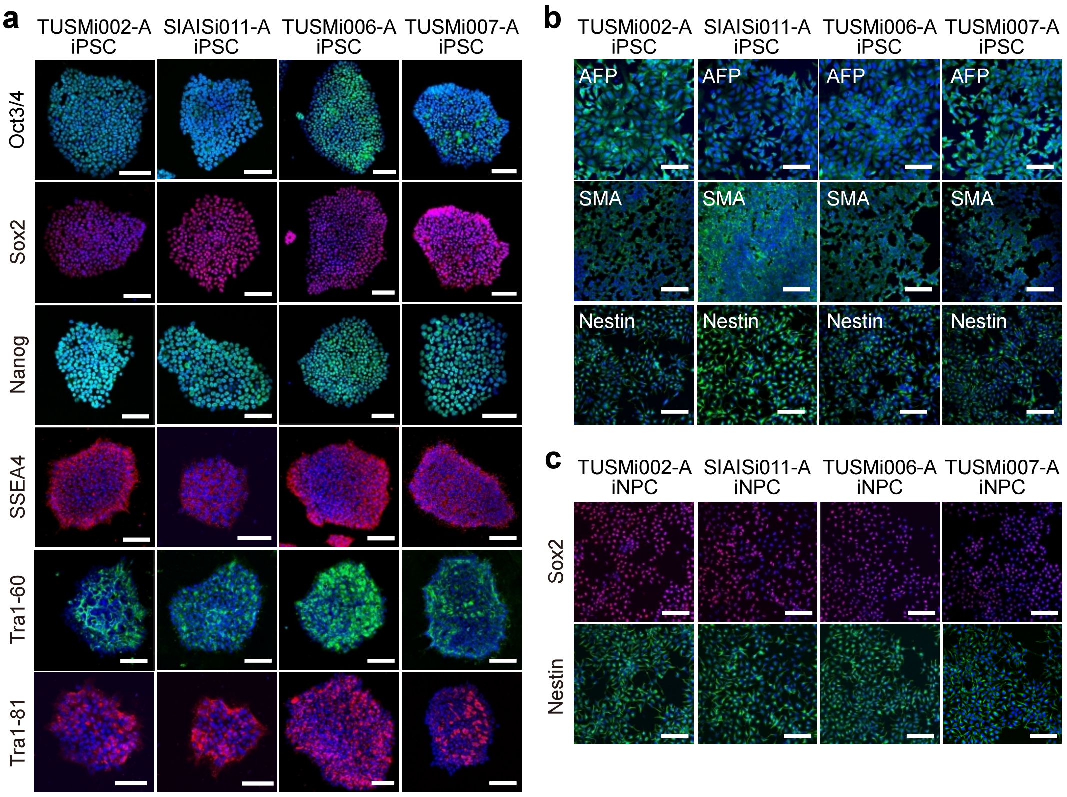
a
TUSMi002-A iPSC
SIAISi011-A iPSC
TUSMi006-A iPSC
TUSMi007-A iPSC
Oct3/4
Sox2
Nanog
SSEA4
Tra1-60
Tra1-81
b
TUSMi002-A iPSC
SIAISi011-A iPSC
TUSMi006-A iPSC
TUSMi007-A iPSC
AFP
SMA
Nestin
c
TUSMi002-A iNPC
SIAISi011-A iNPC
TUSMi006-A iNPC
TUSMi007-A iNPC
Sox2
Nestin

**Figure S2**

**a**

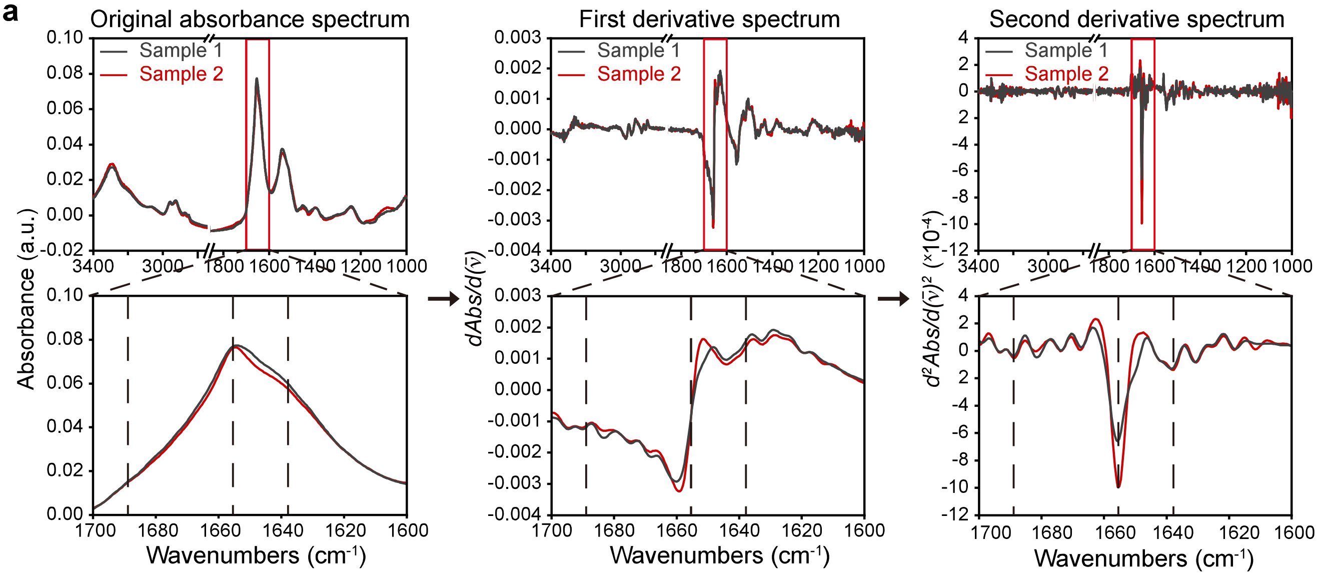


**b**

Infrared spectral absorption bands

| Peak No. | Functional groups | Wavenumber ($cm^{-1}$) | Macromolecules |
|---|---|---|---|
| 1 | Amide A | 3340-3320 | Proteins |
| 2 | Amide B | 3075-3055 | Proteins |
| 3 | Olefinic =CH stretching | 3025-3005 | Fatty acids |
| 4 | $V_{as}(CH_3)$ | 2970-2950 | Fatty acids |
| 5 | $V_{as}(CH_2)$ | 2930-2910 | Fatty acids |
| 6 | $V_s(CH_3)$ | 2880-2860 | Fatty acids |
| 7 | $V_s(CH_2)$ | 2860-2840 | Fatty acids |
| 8 | C=O stretching of lipid esters | 1750-1730 | Fatty acids |
| 9 | C=O antisymmetric stretching: RNA and purine base | 1725-1705 | Nucleic acids |
| 10 | N-H bending of Amide I | 1695-1685 | Proteins |
| 11 | a-helical of Amide I | 1660-1650 | Proteins |
| 12 | β-pleated sheet of Amide I | 1650-1630 | Proteins |
| 13 | C4-C5 and C=N stretching in imidazole ring of DNA | 1620-1600 | Nucleic acids |
| 14 | C4-C5 and C=N stretching in imidazole ring of RNA | 1585-1565 | Nucleic acids |
| 15 | a-helical of Amide II | 1550-1540 | Proteins |
| 16 | β-pleated sheet of Amide II | 1540-1520 | Proteins |
| 17 | “Tyrosine” | 1505-1525 | Proteins |
| 18 | CH2 bending vibrations | 1480-1460 | Fatty acids; Proteins |
| 19 | CH3 bending and CH2 scissoring vibrations | 1460-1450 | Fatty acids; Proteins |
| 20 | COO- symmetric stretching | 1410-1390 | Fatty acids; Proteins |
| 21 | Peptide side chain vibrations | 1320-1300 | Proteins |
| 22 | $Vas(PO^{2-})$ | 1240-1220 | Nucleic acids |
| 23 | CO–O–C antisymmetric stretching vibrations of glycogen and nucleic acid ribose | 1165-1145 | Fatty acids; Nucleic acids |
| 24 | $Vs(PO^{2-})$ | 1090-1070 | Nucleic acids |
| 25 | C-O stretching vibratios:DNA | 1070-1050 | Nucleic acids |
| 26 | C-O stretching vibratios:RNA | 1060-1040 | Nucleic acids |
| 27 | Mainly from glycogen | 1030-1010 | Fatty acids |

### Figure S3

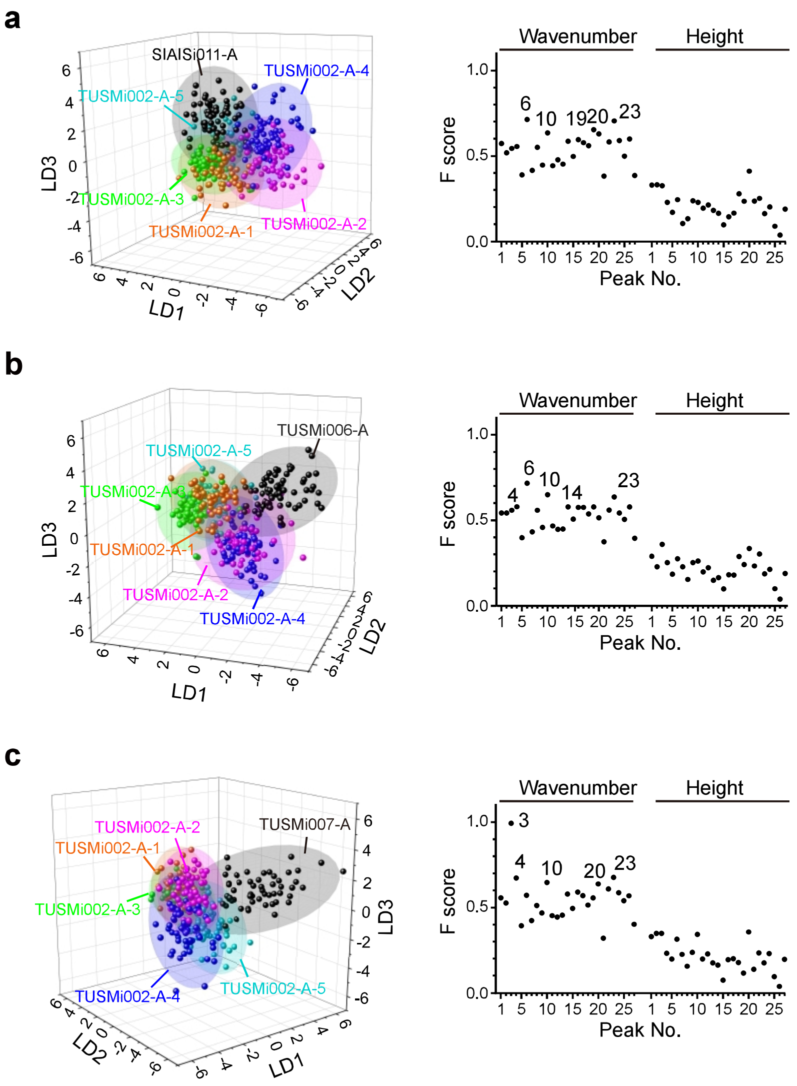
a
SIAISi011-A
TUSMi002-A-4
TUSMi002-A-5
TUSMi002-A-3
TUSMi002-A-1
TUSMi002-A-2
LD1
LD2
LD3
Wavenumber
Height
F score
Peak No.
b
TUSMi006-A
TUSMi002-A-5
TUSMi002-A-3
TUSMi002-A-1
TUSMi002-A-2
TUSMi002-A-4
c
TUSMi002-A-2
TUSMi002-A-1
TUSMi007-A
TUSMi002-A-3
TUSMi002-A-4
TUSMi002-A-5

#### Figure S4

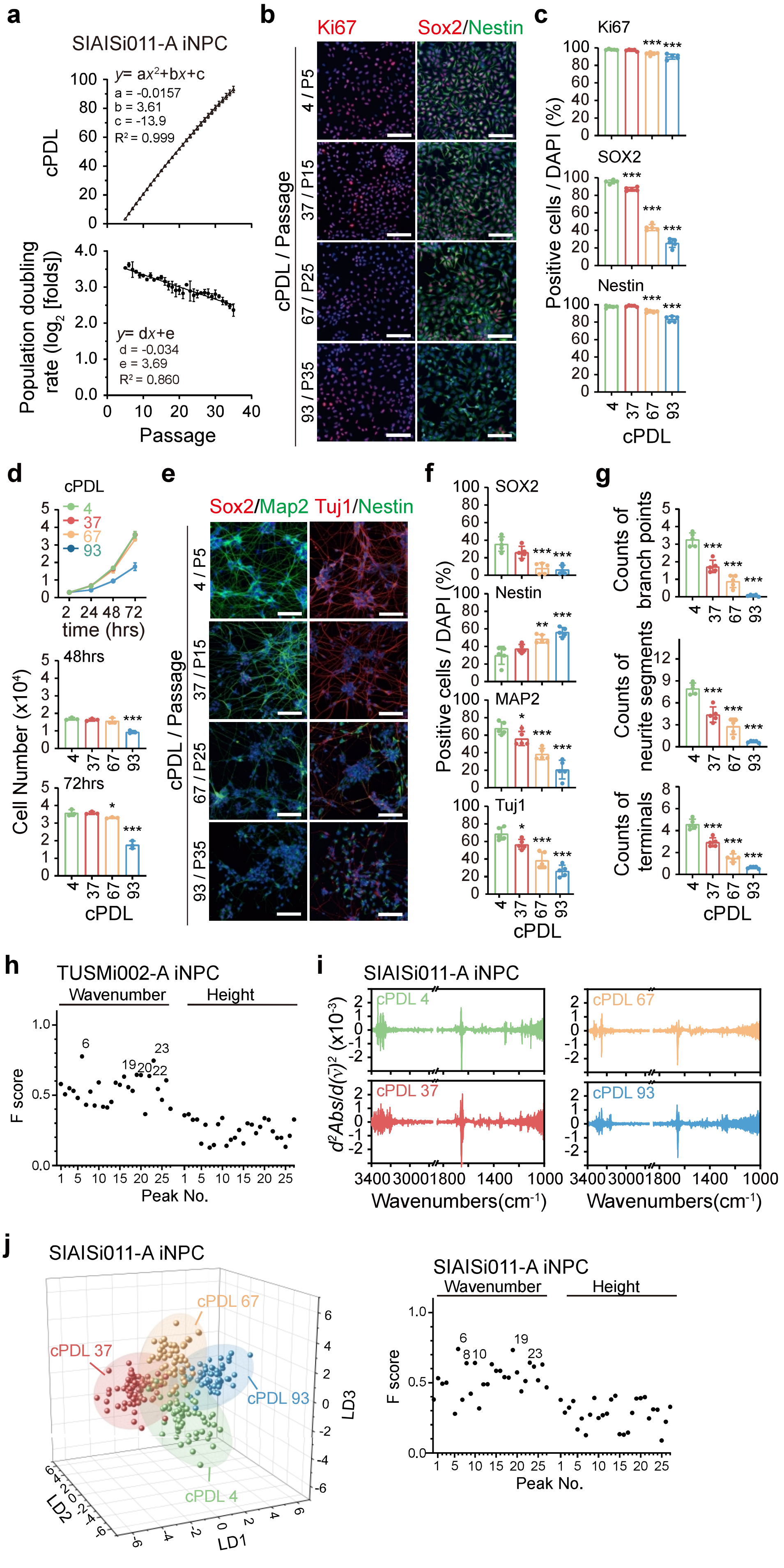

a
SIAISi011-A iNPC
cPDL
y= ax²+bx+c
a = -0.0157
b = 3.61
c = -13.9
R² = 0.999
Population doubling rate (log₂ [folds])
y= dx+e
d = -0.034
e = 3.69
R² = 0.860
Passage
b
Ki67
Sox2/Nestin
cPDL / Passage
4 / P5
37 / P15
67 / P25
93 / P35
c
Ki67
SOX2
Nestin
Positive cells / DAPI (%)
cPDL
d
cPDL
4
37
67
93
time (hrs)
48hrs
72hrs
Cell Number (x10⁴)
cPDL
e
Sox2/Map2 Tuj1/Nestin
cPDL / Passage
4 / P5
37 / P15
67 / P25
93 / P35
f
SOX2
Nestin
MAP2
Tuj1
Positive cells / DAPI (%)
cPDL
g
Counts of branch points
Counts of neurite segments
Counts of terminals
cPDL
h
TUSMi002-A iNPC
Wavenumber
Height
F score
Peak No.
i
SIAISi011-A iNPC
cPDL 4
cPDL 67
cPDL 37
cPDL 93
d²Abs/d(ν)² (x10⁻³)
Wavenumbers(cm⁻¹)
Wavenumbers(cm⁻¹)
j
SIAISi011-A iNPC
cPDL 67
cPDL 37
cPDL 93
cPDL 4
LD1
LD2
LD3
SIAISi011-A iNPC
Wavenumber
Height
F score
Peak No.

**Figure S5**

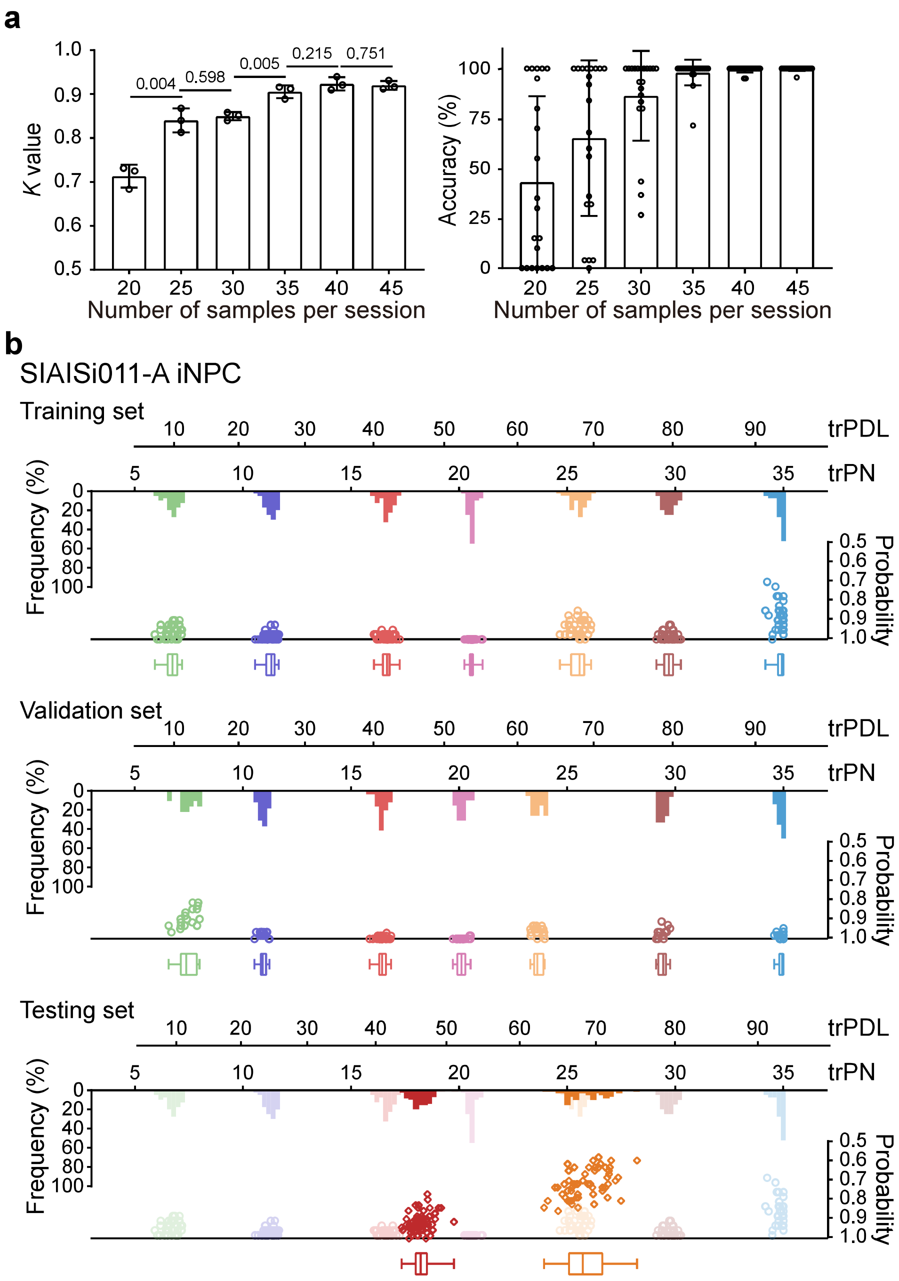


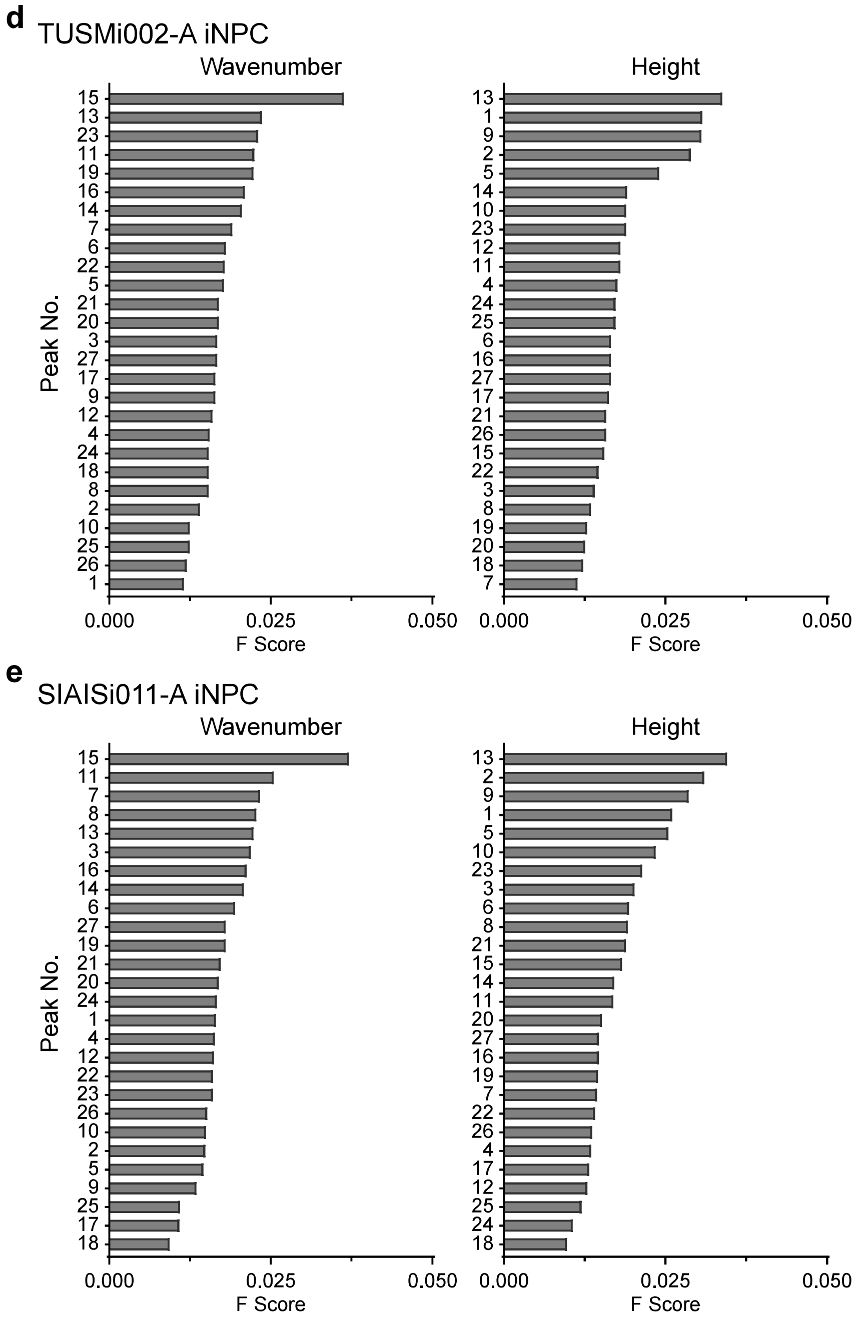


**c**

| | MSE | RMSE | MAE | MAPE | $R^2$ |
|---|---|---|---|---|---|
| Training set | 1.72 | 1.31 | 1.02 | 8.51 | 0.98 |
| Validation set | 1.74 | 1.32 | 1.07 | 8.74 | 0.98 |
| Testing set | 3.52 | 1.88 | 1.66 | 9.39 | 0.82 |

### Figure S6

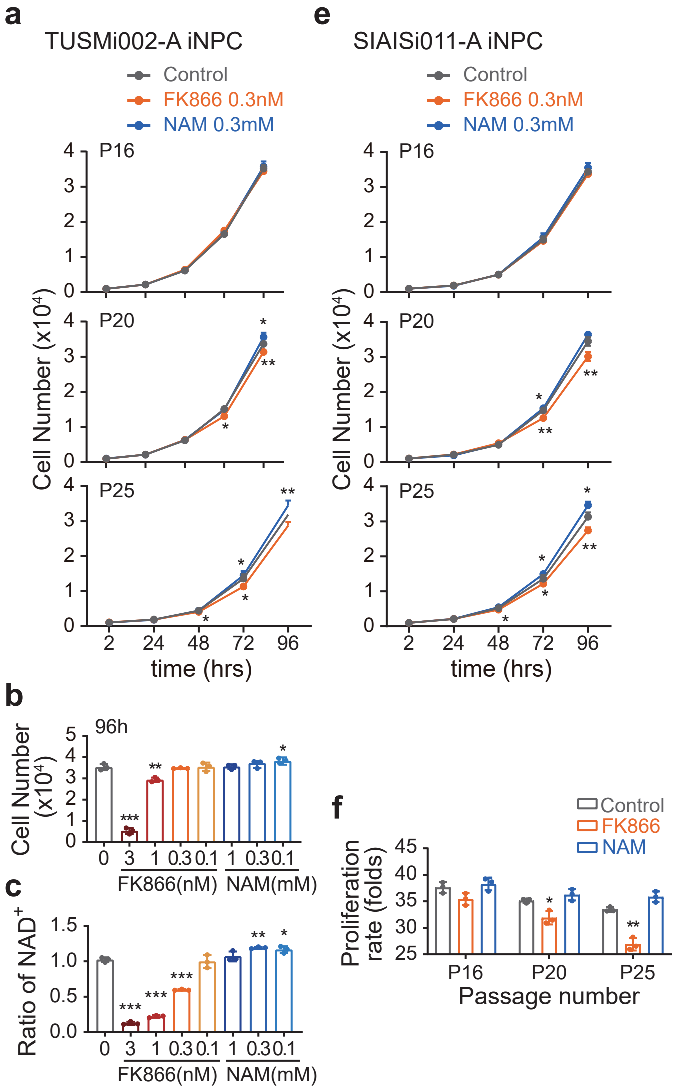


d

| Passage number | SIAISi011-A iNPC | | |
|---|---|---|---|
| | Control | FK866 | NAM |
| P16 | 38.37±1.42 | 38.08±0.11 | 38.40±0.34 |
| P20 | 52.67±1.11 | 52.00±1.36 | 52.98±0.60 |
| P25 | 68.32±0.92 | 67.12±0.68 | 69.17±0.53 |

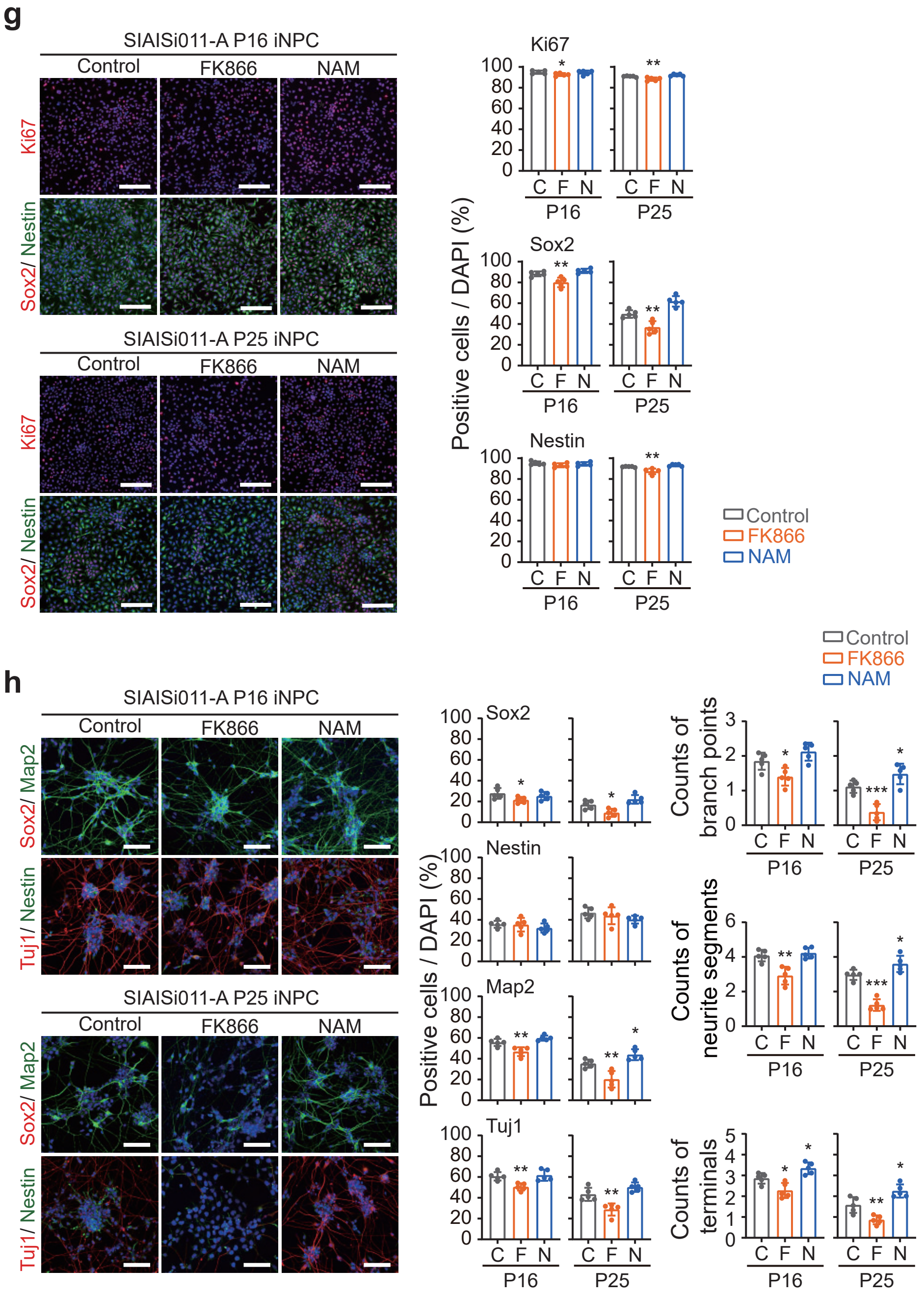

#### Figure S7

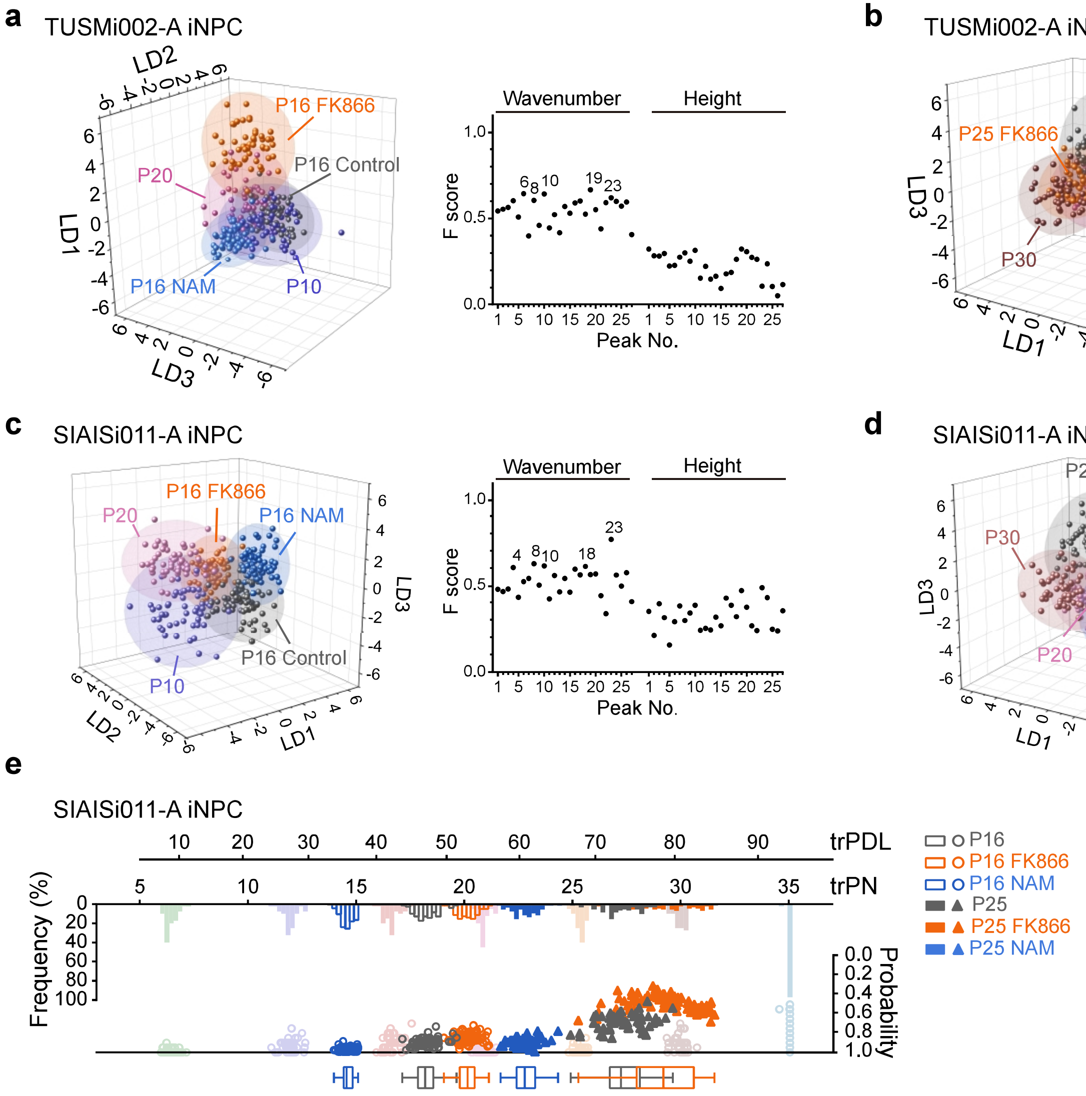
a
TUSMi002-A iNPC
P16 FK866
P16 Control
P20
P16 NAM
P10
LD1
LD2
LD3
Wavenumber
Height
F score
Peak No.
b
TUSMi002-A iNPC
P25 Control
P25 FK866
P25 NAM
P30
P20
c
SIAISi011-A iNPC
P20
P16 FK866
P16 NAM
P16 Control
P10
d
SIAISi011-A iNPC
P25 Control
P30
P25 FK866
P20
P25 NAM
e
SIAISi011-A iNPC
trPDL
trPN
Frequency (%)
Probability
P16
P16 FK866
P16 NAM
P25
P25 FK866
P25 NAM

**Supplement table1: Detailed results of the generated CIADEX model for TUSMi002 - A iNPC and SIAISi011 - A iNPC**

Detailed results for TUSMi002 - A iNPC and SIAISi011-A iNPC across three datasets (training, validation, and testing sets). The median trPN for each population and the corresponding probability, accompanied by the 95% confidence intervals, were listed.

**Supplement table 1**: Detailed results of the CIADEX model

| | | TUSMi002-A iNPC | | | SIAISi011-A iNPC | | |
|---|---|---|---|---|---|---|---|
| | | Training set | Validation set | Testing set | Training set | Validation set | Testing set |
| P5 | trPN | 6.80 (6.72-6.97) | 6.97 (6.78-7.01) | | 7.41 (7.24-7.66) | 7.38 (7.13-7.88) | |
| | probability | 0.98 (0.96-0.98) | 0.96 (0.95-1.00) | | 0.91 (0.88-0.93) | 0.90 (0.84-0.93) | |
| P10 | trPN | 11.81 (11.68-11.90) | 11.93 (11.69-12.16) | | 11.01 (10.92-11.12) | 10.92 (10.82-11.09) | |
| | probability | 0.97 (0.97-0.98) | 0.97 (0.93-0.98) | | 0.98 (0.96-0.98) | 0.98 (0.96-0.98) | |
| P15 | trPN | 16.34 (16.28-16.41) | 16.33 (16.16-16.60) | 16.65 (16.46-16.74) | 16.55 (16.41-16.60) | 16.44 (16.34-16.61) | 18.22 (18.06-18.34) |
| | probability | 0.97 (0.97-0.98) | 0.98 (0.95-0.98) | 0.94 (0.94-0.95) | 1.00 (1.00-1.00) | 1.00 (0.98-1.00) | 0.90 (0.88-0.91) |
| P20 | trPN | 21.38 (21.32-21.47) | 21.30 (21.02-21.45) | | 20.14 (19.98-20.22) | 20.11 (19.92-20.30) | |
| | probability | 0.98 (0.97-1.00) | 0.98 (0.97-0.98) | | 1.00 (1.00-1.00) | 1.00 (1.00-1.00) | |
| P25 | trPN | 25.69 (25.60-25.78) | 25.86 (25.65-26.09) | 23.63 (23.50-23.77) | 23.49 (23.41-23.64) | 23.62 (23.46-23.91) | 25.73 (25.31-26.25) |
| | probability | 0.98 (0.98-1.00) | 0.96 (0.96-1.00) | 0.95 (0.93-0.96) | 0.96 (0.95-0.97) | 0.95 (0.95-0.97) | 0.70 (0.67-0.72) |
| P30 | trPN | 30.06 (29.94-30.15) | 30.11 (30.02-30.26) | | 29.57 (29.48-29.62) | 29.45 (29.21-29.57) | |
| | probability | 0.96 (0.93-0.96) | 0.96 (0.95-0.98) | | 0.98 (0.96-0.98) | 0.96 (0.96-0.98) | |
| P35 | trPN | 35.00 (35.00-35.00) | 35.00 (35.00-35.00) | | 34.98 (34.89-35.00) | 34.94 (34.73-35.00) | |
| | probability | 0.93 (0.90-0.95) | 0.97 (0.92-0.97) | | 0.98 (0.96-0.98) | 0.98 (0.96-1.00) | |

trPN and the corresponding probability are presented as median with 95% confidence intervals.
